# Pre-bound State Discovered in the Unbinding Pathway of Fluorinated Variants of the Trypsin-BPTI Complex Using Random Acceleration Molecular Dynamics Simulations

**DOI:** 10.1101/2024.02.22.581541

**Authors:** Leon Wehrhan, Bettina G. Keller

## Abstract

The serine protease trypsin forms a tightly bound inhibitor complex with Bovine Pancreatic Trypsin Inhibitor (BPTI). The complex is stabilized by the P1 residue Lys15, which interacts with the negatively charged amino acids at the bottom of the S1 pocket. Truncating the P1 residue of wildtype BPTI to *α*-aminobutyric acid (Abu) leaves a complex with moderate inhibitor strength, which is held in place by additional hydrogen bonds at the protein-protein interface. Fluorination of the Abu residue partially restores inhibitor strength. The mechanism with which fluorination can restore the inhibitor strength is unknown and accurate computational investigation requires knowledge of the binding and unbinding pathways. The preferred unbinding pathway is likely to be complex, as encounter states have been described before and unrestrained Umbrella Sampling simulations of these complexes suggest additional energetic minima. Here, we use Random Acceleration Molecular Dynamics to find a new metastable state in the unbinding pathway of Abu-BPTI variants and wildtype BPTI from trypsin, which we call the pre-bound state. The pre-bound state and the fully bound state differ by a substantial shift in the position, a slight shift in the orientation of the the BPTI variants and change in the interaction pattern. Particularly important is the breaking of three hydrogen bonds around Arg17. Fluorination of the P1 residue lowers the energy barrier of the transition between fully bound state and pre-bound state and also lowers the energy minimum of the pre-bound state. While the effect of fluorination is in general difficult to quantify, here it is in part caused by a favorable stabilization of a hydrogen bond between Gln194 and Cys14. The interaction pattern of the pre-bound state offers insight into the inhibitory mechanism of BPTI and might add valuable information for the design serine protease inhibitors.

## Introduction

Proteases are enzymes that play a crucial role in the breakdown of peptides by catalyzing the hydrolysis of peptide bonds. Among them, serine proteases form a subgroup which catalyze this reaction via a serine residue in their active site. Serine proteases are found in most life forms, including bacteria, viruses, fungi, plants, and animals. They play essential roles in digestion, signal transduction, blood clotting, immune responses, and other cellular functions. In humans, serine proteases are important drug targets for many diseases including cardiovascular, cancer and infectious diseases.^1,2^ An example of a serine protease is trypsin, a mammalian digestive enzyme. It has been widely used as model system for serine proteases, since it exhibits the most prevalent fold for proteases in humans and higher organisms. ^3,4^

Protein-protein complexes involving trypsin are stabilized by a long positively charged residue located on the binding protein, the P1 residue, which reaches into the deep S1 binding pocket of trypsin (Schechter and Berger notation).^5^ The S1 pocket is lined with negatively charged residues which either bind directly to the positive charge or via water-mediated contacts.^6,7^ Trypsin’s catalytic site is located at the rim of the S1 binding pocket. Most proteins that bind to trypsin in this manner are cleaved at the C-terminal side of their P1 residue and thus act as substrates. Some proteins, despite binding to the S1 pocket, are not cleaved and instead act as inhibitors towards trypsin. Examples are Bovine Pancreatic Trypsin Inhibitor (BPTI), Antitrypsin or Serpins. Several mechanisms have been put forward to explain why these proteins are not hydrolyzed by trypsin, but instead form such a stable trypsin-inhibitor complex. Possibly, initial hydrolysis might take place, but re-ligation of the cleaved bond is fast and thus favoured over release of the hydrolyzed product.^8^ In the clogged gutter mechanism,^9^ the hydrolyzed products are bound in a tightly and specific orientation to trypsin, such that product release is hindered.

We here focus on BPTI, which is an exceptionally well studied protein^9–13^ and inhibits trypsin with an extraordinarily high binding affinity (the binding constant is *K_i_* = 5 · 10^−24^*M*).^14^ Its P1 residue is Lys15, which forms water mediated bonds to Asp189 and Ser190 at the bottom of the S1 pocket. The importance of Lys15 for the binding process has been demonstrated by kinetic studies with BPTI mutants, where the K15A mutant BPTI shows dramatically decreased binding affinity.^14^ Interestingly, the K15A mutant is also the only variant that has a significantly decreased association rate, highlighting the importance of the P1 residue for the trypsin-BPTI recognition.

While for complexes of proteins with small molecules, like the trypsin-benzamidine complex, the full energy landscape of the binding and unbinding process has been calculated,^15^ the computational characterization of the binding-equilibrium in protein-protein complexes, like the BPTI-trypsin complex, is much more challenging.^16,17^ The reasons for this include the slow movements of macromolecules along translational and rotational degrees of freedom. Also, the number of possible contact conformations of a protein-protein complex far exceeds that of a protein-small-molecule complex. In computational studies of protein-protein complexes additional restraints to the relative position and orientation may be applied to increase the sampling of the binding/unbinding process. ^18,19^ However this requires knowledge of the exact binding/unbinding path to obtain an accurate free energy profile and characterize relevant intermediate states.

Kahler et al. ^20^ studied the binding/unbinding process of wildtype BPTI with trypsin using unbiased simulations, seeded by umbrella simulations. They describe the binding/unbinding process as a two-step mechanism, in which trypsin and BPTI recognize each other first through Coulomb interactions and form encounter states before moving on to form the fully bound protein-protein complex.

We here study the unbinding process of BPTI variants where the Lys15 residue has been mutated to *α*-aminobutyric acid (Abu) and its mono- (MfeGly), di- (DfeGly) and tri-fluorinated (TfeGly) variants. These BPTI variants are not cleaved by trypsin, but instead act as moderate inhibitors with half-maximal inhibitory concentration (IC50) of IC_50_ = 4 · 10^−7^ M (Lys15Abu) and IC_50_ = 6 · 10^−8^ M (Lys15TfeGly).^21,22^

Interestingly, the inhibitor strength of the four BPTI variants systematically increases with increasing fluorination. An initial hypothesis based on crystal structures of the complexes suggested that this increase in binding affinity could be traced back to direct and specific interactions of the fluorine substituents with the water molecules in the S1 pocket. ^21^ In a recent computational study with Molecular Dynamics (MD) simulations, we did not find significant differences in the water structure or water-protein interaction strength across the four variants of the BPTI-trypsin complex,^22^ and thus could not confirm this hypothesis. However, a rough scan using umbrella sampling of the unbinding pathways hinted at a second free-energy minimum, next to the bound state. This pre-bound state was closer to the bound state than encounter states,^20^ which could also be identified in our scan of the unbinding pathway. The existence of a pre-bound state might offer insights into why BPTI acts as a inhibitor rather than a substrate to trypsin, and might open up new avenues for the design of trypsin inhibitors.

In this contribution, we investigate the unbinding path of the four BPTI variants Abu-BPTI, MfeGly-BPTI, DfeGly-BPTI and TfeGly-BPTI using Random Acceleration Molecular Dynamics (RAMD) simulations.^23–26^ Additionally, we also study the unbinding process of wildtype-BPTI.

RAMD is an enhanced sampling method that applies an additional biasing force, which is randomly redirected throughout the simulation, to the center of mass of a ligand and thereby facilitates the exploration of curved unbinding pathways.^23^ RAMD has been used frequently for complexes of proteins with small molecules, but, to the best of our knowledge, this is the first application of RAMD on a protein-protein complex. Our goal is to verify the presence of the pre-bound state and to explain its stability.

## Methods

### Collective Variables

We constructed the collective variables describing the position and orientation of the BPTI variants with respect to trypsin from the positions of three reference points in trypsin (T1, T2, T3) and three reference points in (Abu, MfeGly, DfeGly, TfeGly)-BPTI (B1, B2, B3), adapted from ref. 18. The three reference points for the enzyme trypsin were defined to be the center-of-mass of the backbone of the whole enzyme (T1), the backbone of Val233-Ala241 (T2) and the backbone of Gln46-Leu67 (T3). The reference points of the ligands (Abu, MfeGly, DfeGly, TfeGly)-BPTI were defined to be the center-of-mass of the backbone of the whole ligand (B1), the backbone of Ala48-Thr54 (B2) and the backbone of Cys14-Ala16 (B3). The positions of the reference points in the starting structure are sketched in fig.1. The main collective variable is the center-of-mass distance *r* between the enzyme and the ligand (T1-B1). The angle *θ_o_* (T1-B1-B2) and the dihedrals *ϕ_o_* (T2-T1-B1-B2) and *ψ_o_* (T1-B3-B1-B2) describe the orientation of the ligand with respect to the enzyme. The angle Θ*_p_* (T2-T1-B1) and the dihedral Φ*_p_* (T3-T2-T1-B1) describe the position of the ligand with respect to the enzyme. The collective variables *r*, Θ*_p_*, Φ*_p_*, *θ_o_*, *ϕ_o_* and *ψ_o_* were calculated with Plumed^27,28^ 2.8 for all simulations.

**Figure 1:**
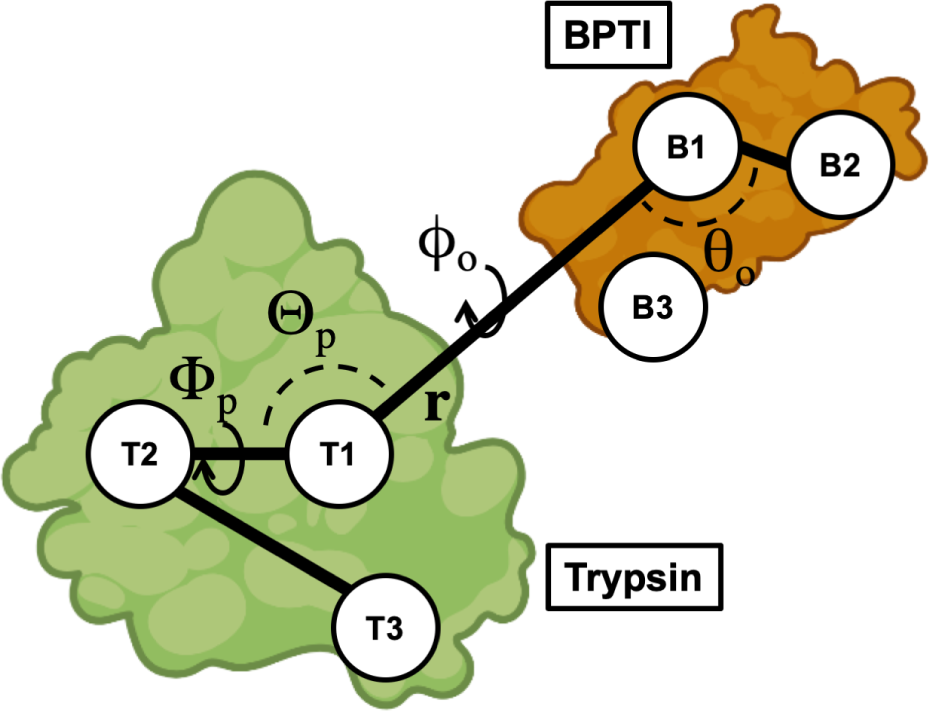
Method to construct collective variables that describe position and orientation of the BPTI variants with respect to trypsin. The position relative to trypsin is described by Θ*_p_* (T2-T1-B1) and Φ*_p_* (T3-T2-T1-B1). The orientation is described by *θ_o_* (T1-B1-B2), *ϕ_o_* (T2-T1-B1-B2) and *ψ_o_* (T1-B3-B1-B2). **r** (T1-B1) is the center-of-mass distance. Compare ref 18.

### Molecular Dynamics General Methods

We ran all Molecular Dynamics (MD) simulations using Gromacs ^29–31^ software and our self-parameterized Amber14SB force field.^22,32–34^ Energy minimizations were conducted with the steepest descend algorithm. Equilibrations in the NVT ensemble were using a velocity rescaling scheme with a stochastic term^35^ to keep the temperature at 300 K and harmonic restraints were applied on all protein heavy atom positions. Subsequent equilibrations in the NPT ensemble without restraints made use of the same velocity rescaling scheme with a stochastic term and the Parinello-Rahman barostat^36^ to keep the temperature at 300 K and the pressure at 1.0 bar. Production MD simulations were run in the NPT ensemble at 300 K and 1.0 bar using the same thermostat and barostat. All MD simulations were performed with the leap-frog integrator and an integration time step of 2 fs. Bond lengths involving hydrogen atoms were kept constant using the LINCS^37^ algorithm. Long range electrostatic interactions above a cutoff distance of 1.0 nm were treated using the PME ^38^ algorithm.

### Starting Structure Preparation

Starting structures were generated from the crystal structure of the TfeGly-BPTI-trypsin complex (pdb code: 4Y11).^21^ Co-solutes and ions were deleted and appropriate hydrogen atoms were added to the crystal structure using the pdbfixer software. The three histidine side chains in the complex were protonated at N(*ɛ*) and N(*δ*). From this initial starting structure, the TfeGly residue was transformed into DfeGly, MfeGly and Abu, respectively, to yield one initial starting structure for every BPTI variant. For the RAMD simulations, the initial starting structures were placed inside a cubic box with periodic boundary conditions with 2.1 nm distance between the solute and the box edges and solvated in TIP3P^39^ water. The systems were energy minimized and equilibrated in the NVT ensemble for 100 ps, followed by an equilibration in the NPT ensemble for 1 ns. To generate two more replicas for each of the complexes, two subsequent simulations of 10 ns were run with the equilibrated starting structures to yield the starting structures for the next replicas.

### Random Acceleration Molecular Dynamics (RAMD)

We used Random Acceleration Molecular Dynamics (RAMD) to explore unbinding pathways of (Abu, MfeGly, DfeGly, TfeGly)-BPTI and wildtype BPTI from trypsin using Gromacs2020.5-RAMD-2.0. Two pull groups were defined: one included all atoms of trypsin, the other included all atoms of (Abu, MfeGly, DfeGly, TfeGly)-BPTI. A random force acting between the two pull groups with a magnitude of 3500 kJ/(mol · nm) was applied. The appropriate force was estimated by running single RAMD simulations of the TfeGly-BPTI-trypsin complex starting with a force of 250 kJ/(mol · nm) and raising the force by 250 kJ/(mol · nm) every simulation until dissociation within 10 ns was achieved. Retrospectively, higher forces between 4000 kJ/(mol · nm) and 5500 kJ/(mol · nm) were tested with the same system to see when the RAMD simulations would fail to detect the pre-bound state at all. Three starting structures for each of the four complexes of trypsin with Abu-BPTI, MfeGly-BPTI, DfeGly-BPTI and TfeGly-BPTI were generated as described above. For every of these replicas, ten RAMD simulations were run from the same starting structure, where the random seed of the random force was changed. The simulations were stopped after dissociation was achieved, the maximum length of the simulations was set to be 40 ns. In the beginning of the simulations, the direction of the biasing force was chosen at random. Throughout the simulations, after every 100 fs, the direction of the force was either retained, if the center of mass of the second pull group moved by more than 0.0025 nm, or changed randomly, if this was not the case. Snapshots were extracted every 2 ps.

### Unbiased Molecular Dynamics of Pre-bound state

We ran unbiased Molecular Dynamics simulations to sample the fully bound state and the pre-bound state of the four complexes of trypsin with Abu-BPTI, MfeGly-BPTI, DfeGly-BPTI, TfeGly-BPTI and wildtype BPTI using Gromacs2021.5,^29–31^ patched with Plumed^27,28^ 2.8. For every complex and state, 20 simulations of 50 ns length were run, totalling 160 simulations with an aggregated length of 8 *µ*s. Initial starting structures for the simulations of the fully bound state were generated from the pdb structure of TfeGly-BPTI as described above. The initial starting structures were placed in a cubic box with periodic boundary conditions with 1.5 nm distance between the solute and the box edges and solvated in TIP3P water. Then, 20 starting structures of every complex for the production MD simulations were generated by individually energy minimizing the systems, followed by equilibration in the NVT ensemble for 100 ps and equilibration in the NPT ensemble for 1 ns. Initial starting structures for the simulations of the pre-bound state were generated by extracting the coordinates of all protein atoms of 20 snapshots of one RAMD simulation of the TfeGly-BPTI-trypsin complex, when the system was in the pre-bound state. The TfeGly residue was transformed into DfeGly, MfeGly and Abu to yield initial starting structures for the other three complexes. The initial starting structures were individually placed in a box with periodic boundary conditions and energy minimized and equilibrated in the same way as the starting structures for the fully bound state. Production MD simulations were run for a length of 50 ns for every replica. Snapshots were extracted every 10 ps.

### Analysis of Distances, Hydrogen Bonds and SASA

We calculated atomic distances and detected hydrogen bonds in simulation snapshots using the python package MDTraj^40^-1.9.4. Hydrogen bonds were detected using the Wernet-Nilsson criterion^41^ implemented in MDTraj:

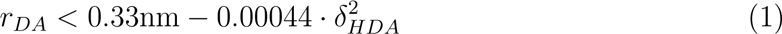

with the donor-acceptor distance *r_DA_* and the angle between hydrogen atom, donor and acceptor *δ_HDA_*.

Distances to the nitrogens in the guanidine moieties of arginine side chains were calculated by computing the distance of the respective interaction partner to all three nitrogen atoms of guanidine moiety and taking the minimum of these three distances for every simulation snapshot.

Solvent Accessible Surface Area (SASA) was calculated using the MDTraj implementation of the Shrake Rupley algorithm.^42^ The SASA for residues was calculated by summing over the atoms in each residue.

### Umbrella Sampling

We conducted Umbrella Sampling using Gromacs2021.5,^29–31^ patched with Plumed^27,28^ 2.8, based on the distance between the backbone oxygen of Phe41 (Phe41-O) of trypsin and the backbone nitrogen of Arg17 (Arg17-N) of the BPTI variants as the main collective variable *ξ*. Starting structures for the umbrella windows were generated starting from the fully bound crystal structure of the Abu-BPTI-trypsin complex as described above. An initial harmonic restraint with a force constant of 6276 kJ/(mol· nm^2^) was placed at *ξ* = 0.25 nm. The system was equilibrated in the NPT ensemble with this harmonic restraint for 500 ps to yield the starting structure of the first umbrella window. Then, the harmonic potential was shifted by 0.05 nm and a new NPT equilibration of 500 ps was run to yield the starting structure of the next window. This procedure was repeated until *ξ* reached a value of 0.70 nm. Additional windows were added in-between at *ξ* values of 3.75 nm, 4.25 nm and 4.75 nm to achieve better sampling of the region of the free energy barrier. Finally there were 13 umbrella windows at the following positions of *ξ* (all in nm): 0.250, 0.300, 0.350, 0.375, 0.400, 0.425, 0.450, 0.475, 0.500, 0.550, 0.600, 0.650, 0.700. In each of the umbrella windows, a production MD simulation with a harmonic restraint with a force constant of 6276 kJ/(mol · nm^2^) was run for a length of 30 ns. The potential of mean force profiles were calculated using binless WHAM.^43,44^ Statistical uncertainty was estimated using a simplified Bootstrapping scheme: The simulations of every window were separated into five parts of 6 ns length. Then, for every window, five combinations of four of these parts were constructed by combining all parts but one. The WHAM calculation was performed on all of these five combinations and the mean and standard deviation of the resulting potential of mean force profiles was calculated.

## Results and Discussion

### Protein-Protein Complex between Trypsin and BPTI Variants

We consider BPTI-variants in which the P1 residue is substituted by *α*-aminobutyric acid (Abu) and its mono- (MfeGly), di- (DfeGly) and tri-fluorinated (TfeGly), i.e., K15Abu, K15MfeGly, K15DfeGly, K15TfeGly). The crystal structures of all four BPTI-variants (pdb codes: 4Y0Z, 7PH1, 4Y10, 4Y11, 4Y0Y) are similar to each other and to the wildtype complex.^21,22^ The interaction strength and pattern between the P1 residue and the S1 pocket and water molecules within the S1 pocket did not differ significantly across BPTI variants, and thus did not explain the observed differences in the stability of the protein-protein complexes.^22^ Fig. 2 shows the complex and binding interface of the TfeGly-BPTI complex as a representative.

**Figure 2:**
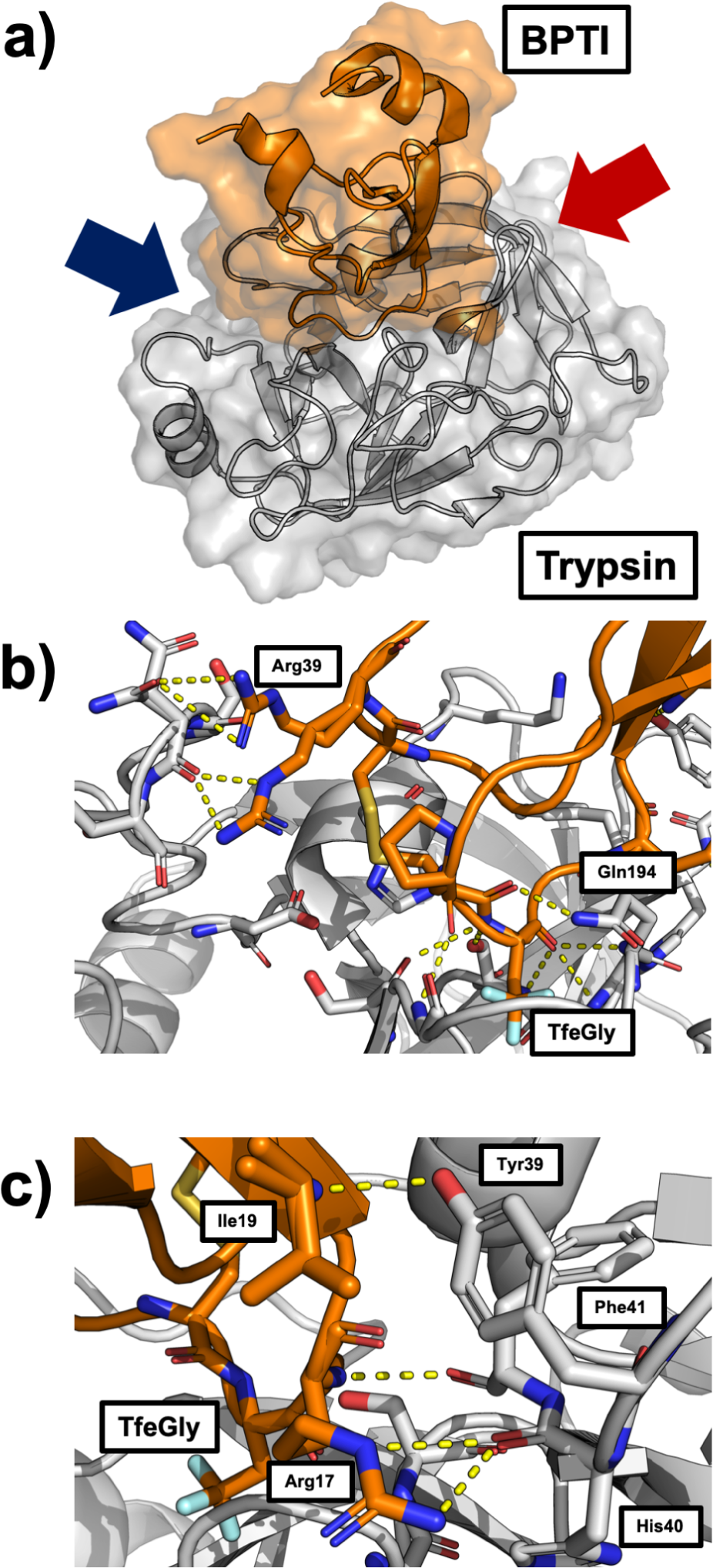
(a) TfeGly-BPTI-trypsin complex with surface and cartoon representation (pdb code: 4Y11). (b) Protein-protein interface of the complex seen from the perspective of the blue arrow. The interactions around the S1 pocket are to the bottom right and the interactions of Arg39 are on the top left. (c) Protein-protein interface of the complex seen from the red arrow. The interactions of Arg17 (P2’) and Ile19 (P4’) are shown.

Besides the S1-P1 interactions, the complex is stabilized by hydrogen bonds throughout the whole protein-protein interface. Fig.2b shows that the P1 residue TfeGly is held in place by seven hydrogen bond like contacts, most notably three backbone interactions holding the backbone carbonyl of the P1 residue in the oxyanion hole of the catalytic pocket. To the left in fig.2b, Arg39 of the BPTI variant can be seen in two alternative conformations, forming interactions to either the side chain or the backbone of Asn97 in trypsin.

Fig. 2c shows the other side of the interface. On this side of the interface, Arg17 (P2’ residue) of the BPTI variant forms interactions with its side chain to the backbone of His40 and with its backbone to the backbone of Phe41. The P4’ residue Ile19 forms an interaction to the side chain of Tyr39 in Trypsin. This interaction has been described as important for the binding of BPTI to Trypsin, as Y39A mutants of Trypsin are less sensitive to BPTI.^3^

### Collective Variables and Metadynamics with Restraints

To investigate the dissociation of the complexes between the BPTI variants and trypsin, we designed a set of collective variables, that describe the position and orientation of the (Abu, Mfegly, DfeGly, TfeGly)-BPTI with respect to trypsin, following Ref. 18. The collective variables are based on the backbone center of mass of the BPTI variants (B1) and trypsin (T1) and two additional points for each of the proteins (T2,T3 and B2, B3), defined as centers-of-mass of well-structured regions inside the proteins (fig. 1). The position of the BPTI-variant relative to trypsin is then given by the distance *r* between B1 and T1, the angle Θ*_p_* = *^̸^* T2-T1-B1, and the dihedral angle Φ*_p_* = *^̸^* T3-T2-T1-B1. The orientation of the BPTI-variant relative to trypsin is given by the angle *θ_o_* = *^̸^* T1-B1-B2 and the dihedrals *ϕ_o_* = *^̸^* T2-T1-B1-B2 and *ψ_o_* = *^̸^* T1-B3-B1-B2.

In an initial attempt to achieve a free energy surface of the binding and unbinding process of (Abu,TfeGly)-BPTI from trypsin, we performed restrained Metadynamics simulations, where we chose the center-of-mass distance between (Abu, TfeGly)-BPTI and trypsin as the main collective variable and used harmonic restraints to restrain the other collective variables to the values of the fully bound complex, which we extracted from the X-ray crystal structure of the TfeGly-BPTI-trypsin complex (pdb code: 4Y11). Our efforts did not yield sufficient sampling of the binding and unbinding process, as after a single unbinding event, the ligand did not find back into the fully bound complex throughout 500 ns Metadynamics simulations, although their orientation and movement around the receptor was restrained (see fig. S1). We conclude, that the preferred binding and unbinding pathway has to be more complex than a simple movement on a straight line defined only by the center-of-mass distance, and likely contains intermediate states.

### Random Acceleration Molecular Dynamics (RAMD)

To study the unbinding pathways of (Abu, MfeGly, DfeGly, TfeGly)-BPTI and wildtype BPTI from trypsin, we performed Random Acceleration Molecular Dynamics (RAMD)^23,24,45^ simulations. RAMD is an enhanced sampling method that applies an additional biasing force to the center-of-mass of a ligand in an otherwise unbiased MD simulation.^23^ If the unbinding process does not make progress despite the biasing force, the direction is of the this force is reoriented in a random direction at regular time intervals. The method was originally invented to discover unbinding pathways of buried protein ligands.^26^

For every of the four Abu-BPTI-variants and wildtype BPTI, we generated three different starting structures and ran 10 simulations with a maximum length of 40 ns for each of these replicas. To achieve dissociation, we needed a force with a high magnitude of 3500 kJ/(mol · nm), which is about an order of magnitude higher than for protein-small molecule systems like benzamidine-trypsin.^23,24^ This might be expected, as according to inhibition assays,^22^ our systems have a binding affinity of −37 kJ/mol to −41 kJ/mol, while benzamidine binds to trypsin with a binding affinity of −22 kJ/mol to −26 kJ/mol.^15^ Moreover, as the complex is held in place by many hydrogen bonds, it is likely that some, if not most, of them have to be broken in a concerted way to achieve dissociation, which would result in a very steep free energy barrier, requiring a strong force to drive the system out of the bound state. Possibly, proteins in general need a higher force constant to be dissociated efficiently compared to small molecules.

For the TfeGly-BPTI-Trypsin complex, fig.3 shows the timeseries of the center-of-mass distance (*r*) as moving average with a moving window of 200 ps. The panels correspond to the three different starting structures, and we show the timeseries of the 10 simulations per starting structure in different colors. See SI (figs. S2-S6) for the corresponding timeseries of DfeGly-, MfeGly-, Abu-, and wildtype BPTI. The timeseries in fig. 3 first fluctuate around the center-of-mass distance of the fully bound complex at around 2.65 nm. Then they tend to transition to a state in which the center-of-mass distance fluctuates between 2.75 nm and 3.00 nm. The systems tends to remain in this state for tens of nanoseconds until they dissociate very rapidly. We call this intermediate state of the protein-protein complex the pre-bound state. We distinguish it from the fully bound state at 2.65 nm center-of-mass distance and from encounter states, which were investigated by Kahler et al. ^20^ and which would lie at center-of-mass distances around 3.00 nm. ^22^

**Figure 3:**
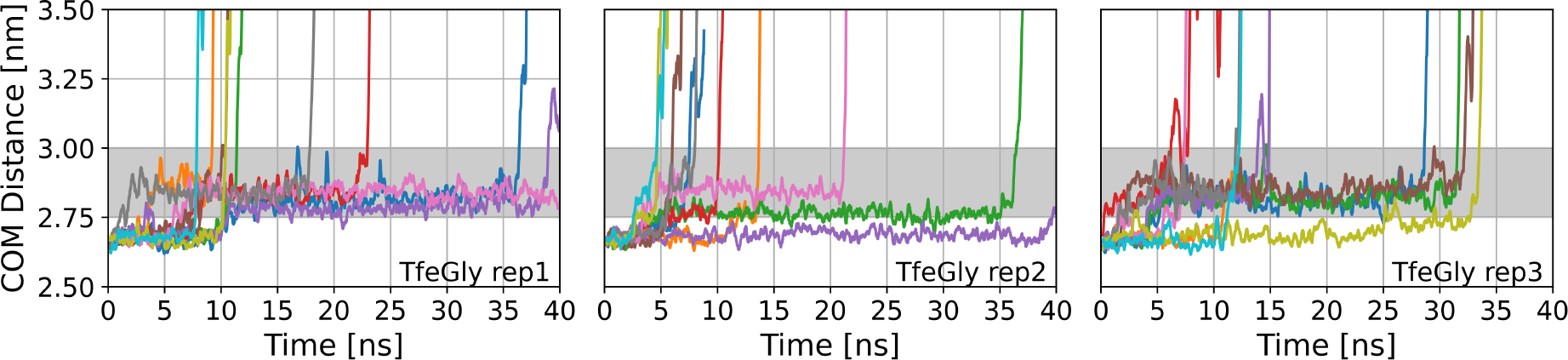
RAMD dissociation timeseries of TfeGly-BPTI. COM = center-of-mass. Grey area shows the center of mass of the pre-bound state. Left panel: replica 1, middle panel: replica 2, right panel:replica 3. 10 RAMD runs per replica.

The pre-bound state occurs in dissociation trajectories of all the four complexes of trypsin with (Abu, MfeGly, DfeGly, TfeGly)-BPTI as well as in the dissociation trajectories of wildtype-BPTI. While in some of the simulations dissociation occurs without visiting the pre-bound state, we observe that in more than half of the trajectories for all BPTI variants, the moving average of the center-of-mass distance remains at least 1 ns between 2.75 nm and 3.00 nm, i.e. the pre-bound state is visited. Some trajectories did not dissociate after 40 ns of RAMD simulation, with some simulations ending in the pre-bound state and others ending in the fully bound state (see SI (tab. S1). The stability of the pre-bound state is remarkable, since throughout the RAMD simulations a strong biasing force designed to dissociate the protein-protein complex acts on the center of mass of the BPTI variant.

Once the system leaves the pre-bound state towards larger center-of-mass distances, the protein-protein complex rapidly dissociates. That is, we do not observe encounter complexes around or above 3.00 nm center of mass distance for any of the BPTI variants in our RAMD simulations. Encounter complexes are typically only weakly bound, and we assume that because of the strong biasing force encounter complexes rapidly dissociated in the RAMD simulations.

Inspecting the RAMD trajectories more closely, we find that the pre-bound state is not only characterized by an increase in center-of-mass distance *r*, but also by a significant shift of the system in the positional collective variables Θ*_p_* and Φ*_p_* compared to the fully bound state (see fig. S7). Fig. 4 shows that the positional variables Θ*_p_* or Φ*_p_* change along with the center-of-mass distance *r* when transitioning from the fully bound state to the pre-bound state. In the orientational variables, *θ_o_*, *ϕ_o_*, *ψ_o_*, we do not find such a correlation, except for a slight shift in the *θ_o_* angle (see. fig. S8).

**Figure 4:**
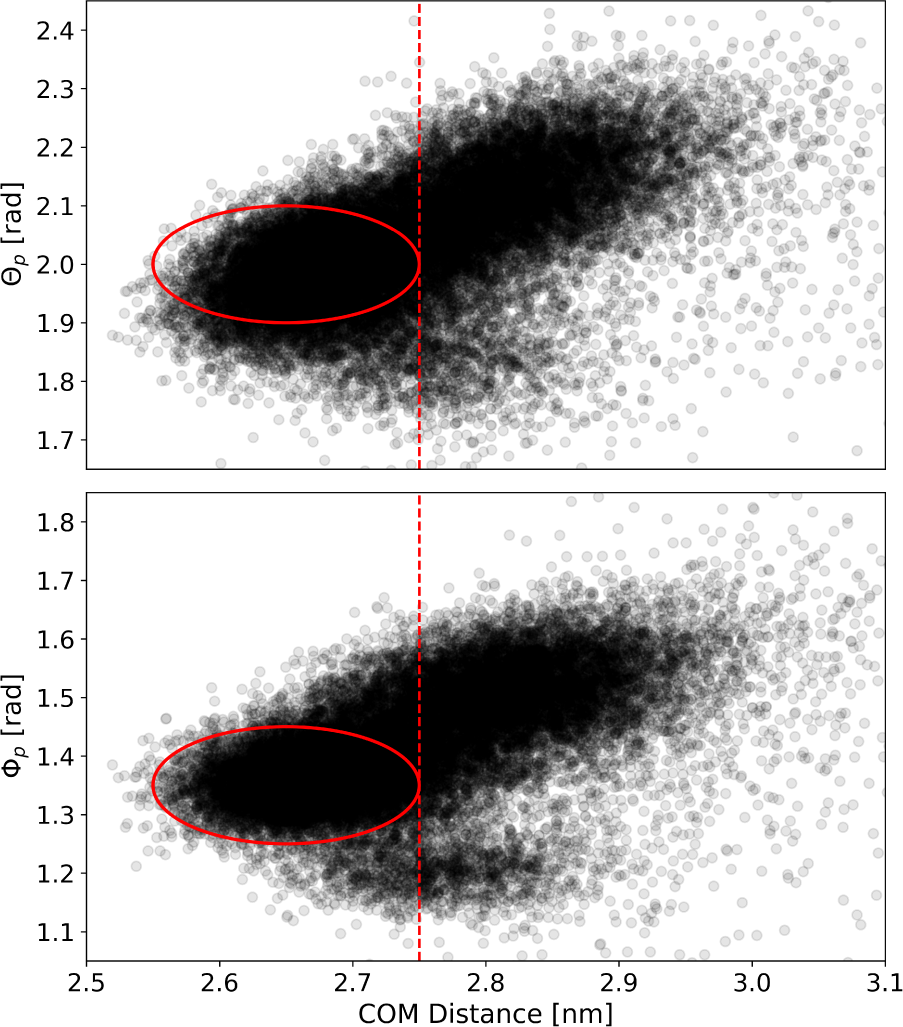
Scatter plots of the center of mass distance and Θ*_p_* (top) and Φ*_p_* (bottom) for all combined RAMD simulations. The red circle marks the position of the fully bound state.

The stability in presence of the biasing force and the systematic change in the positional variables indicate that the pre-bound state might be a chemically relevant state, which is stabilized by different interactions than the fully bound state and separated by a free-energy barrier from the fully bound state. As the fully bound state is held in place tightly by a number of hydrogen bond like interactions, it is likely that some of these interactions must be broken so the pre-bound state can be reached. In the fully bound state of the complex between Trypsin and (Abu, MfeGly, DfeGly, TfeGly)-BPTI there are twelve hydrogen bond like contacts, that can be found by visually inspecting the crystal structures and which are shown in fig. 2. We calculated the frequency with which these interactions are formed as a function of the center-of-mass distance and present the histograms in fig. 5. The criterion for an interaction being in place was a heavy atom distance of less than 0.35 nm. Note that the histograms were generated from RAMD simulations, i.e. non-equilibrium simulations, and therefore do not represent equilibrium distributions.

**Figure 5:**
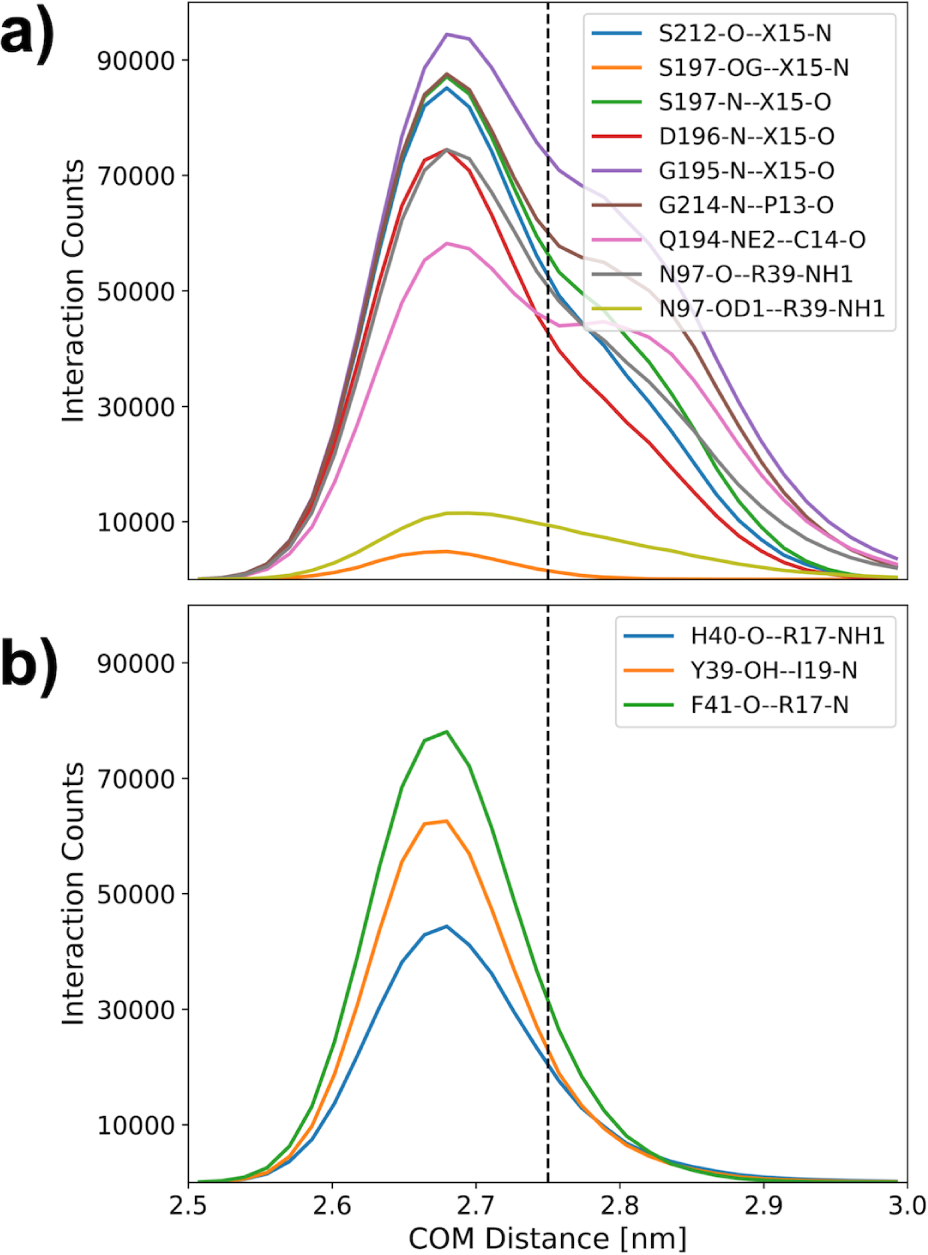
Interaction histogram along center of mass distance in RAMD simulations of the Abu-BPTI variants and trypsin. The criterion for an interaction to be in place was that the involved heavy atoms were separated by a distance less than 0.35 nm. The histograms were generated from the biased (non-equilibrium) RAMD simulations and therefore do not represent a Boltzmann distribution.

One of the twelve interactions is only rarely populated in the fully bound state and not populated at all in the pre-bound state: the hydrogen bond between the side chain hydroxyl oxygens of Serine 197 (Ser197-OG) in trypsin and the amide hydrogen in the backbone of the P1 residue in the BPTI variants (X15-N), shown in orange in fig. 5.a. But since the overall change is small, this interaction is not suited to further define the pre-bound state. Also in fig. 5.a, we show the histograms of eight further interactions, which are present in the fully bound state as well as in the pre-bound state. Their population decreases with increasing center-of-mass distance, but since the change is gradual and there is still significant population in the interval 2.75 nm < *r* < 3.00 nm, it is not plausible that this change in population constitutes a clear free-energy barrier between the fully bound state and the pre-bound state.

Fig. 5.b shows the histogram of three interactions which are highly populated in the fully bound state but rarely populated in the interval 2.75 nm < *r* < 3.00 nm. These are the backbone-backbone interaction between Phe41 and Arg17, the interaction of the backbone of His40 with the side chain of Arg17 and the interaction between the side chain of Tyr39-OH and the backbone of Ile19. (compare fig.2c). The breaking of these three interactions likely contributes to the free energy barrier between the fully bound state and the pre-bound state.

Considering that we used a very high random force of 3500 kJ/(mol · nm), we note that it is remarkable that the systems remain in the pre-bound state for a substantial amount of simulation time, despite the strong bias force introduced to the simulation. To retrospectively test the limits of this method, we ran sets of simulations with the TfeGly-BPTI variant where we increased the magnitude of the random force to even higher values up to 5500 kJ/(mol · nm). The trajectories are shown in fig. S9. We still observe the dissociating system to briefly stay in the region of *r* typical for the pre-bound state for some trajectories with a random force of 5000 kJ/(mol · nm), but not with 5500 kJ/(mol · nm). Hence, we conclude that a random force of 5000 kJ/(mol · nm) is the limit to observe the pre-bound state for this system.

### Unbiased Simulation of the Pre-Bound and Fully Bound state

To further characterize the difference between the fully bound state and the pre-bound state, we ran 20 unbiased simulations of 50 ns each (i.e. 1 *µ*s total simulation time) of the fully bound state and of the pre-bound state in all of the four complexes between trypsin and (Abu, MfeGly, DfeGly, TfeGly)-BPTI and wildtype BPTI. The starting structures for the fully bound state were generated from the crystal structure and the starting structures for the simulations of the pre-bound state were generated from snapshots of the system in the pre-bound state from the RAMD simulations.

The timeseries of the center-of-mass distance *r* for all of the unbiased MD simulations can be found in the SI (fig. S10 for the Abu-BPTI variants and fig. S11 for wildtype-BPTI). With very few exceptions, the systems remained in their starting state throughout the whole simulation time. This indicates that both the fully bound state and the pre-bound state are stable on the timescale of 50 ns.

Fig. 6 compares the equilibrium distributions of the positional variables, *r*, Θ*_p_*, Φ*_p_* of the fully bound state and of the pre-bound state. All BPTI variants have similar distributions (different colors in fig. 6), with the exception of the distributions of Θ*_p_* of wildtype-BPTI, which is shifted towards higher values, compared to the Abu-BPTI variants. However, the distribution differ significantly between the fully bound state and the pre-bound state (solid vs. dashed lines in in fig. 6). In the fully bound state, the systems adopt an average center-of-mass distance of 2.65 nm with a standard deviation of 0.04 nm, while in the pre-bound state the center-of-mass distance *r* amounts to a mean of 2.85 nm with a standard deviation of 0.05 nm. Likewise, the coordinates Θ*_p_* and Φ*_p_* shift to larger values in the pre-bound state. In all three positional coordinates, there is little overlap between the distributions of the fully bound state and the pre-bound state, confirming that the positions that the BPTI variants can occupy in these two states are distinct.

**Figure 6:**
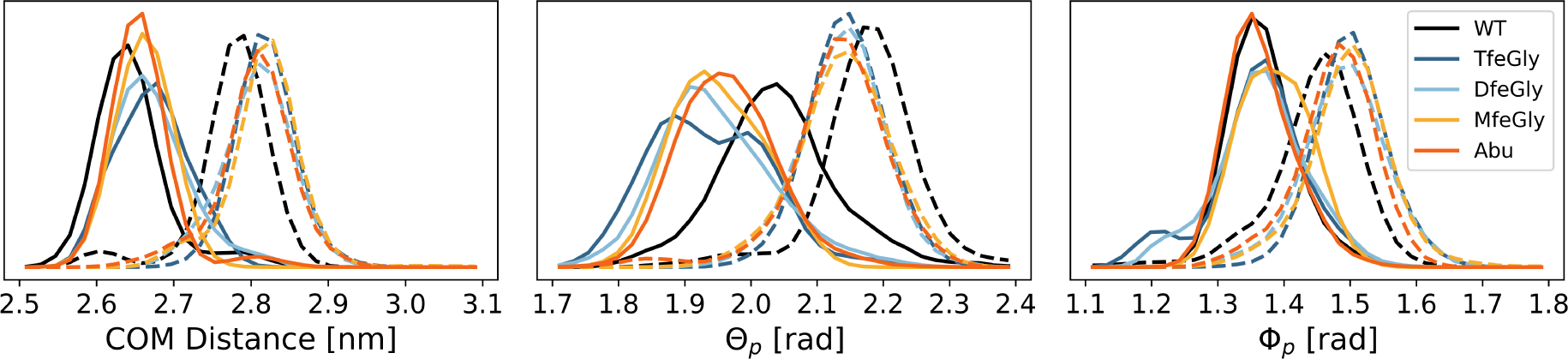
Position of the BPTI variants in the unbiased simulations of the fully bound state (solid lines) and the pre-bound state (dashed lines) as described by the center of mass distance *r* (left), Θ*_p_* (center) and Φ*_p_* (right).WT = wildtype.

The distributions of the orientational variabiles, *θ_o_*, *ϕ_o_*, *ψ_o_*, are included in the SI (fig. S12). In each of the three variables we observe a systematic shift from the distributions of the fully bound state and to those of the pre-bound state, which is most pronounced for *θ_o_*. However, the overlap between the fully bound state distributions and the pre-bound state distributions is larger than for the positional variables. This indicates that the BPTI variant does not gain (much) orientational freedom when transitioning from the fully bound state to the pre-bound state. We provide example snapshots from the unbiased simulations of the fully bound state and the pre-bound state for all of the four BPTI variants in the Supplementary Material.

Fig. 7 shows the relative population of the all hydrogen bonds between the two proteins with at least 0.1 relative population. For this analysis we merged the trajectories of the fully bound state of all four Abu-BPTI variants, and we merged the trajectories of the pre-bound state of all four Abu-BPTI variants (fig. 7a). At this point this is justified, because the twelve interactions do not involve the side chain of the P1 residue and because we did not observe any significant difference in the positional and orientational variables across the four systems (fig. S13). We employed the Wernet-Nilsson criterion^41^ in the MDtraj implementation to identify hydrogen bonds between Trypsin and the BPTI variants.

**Figure 7:**
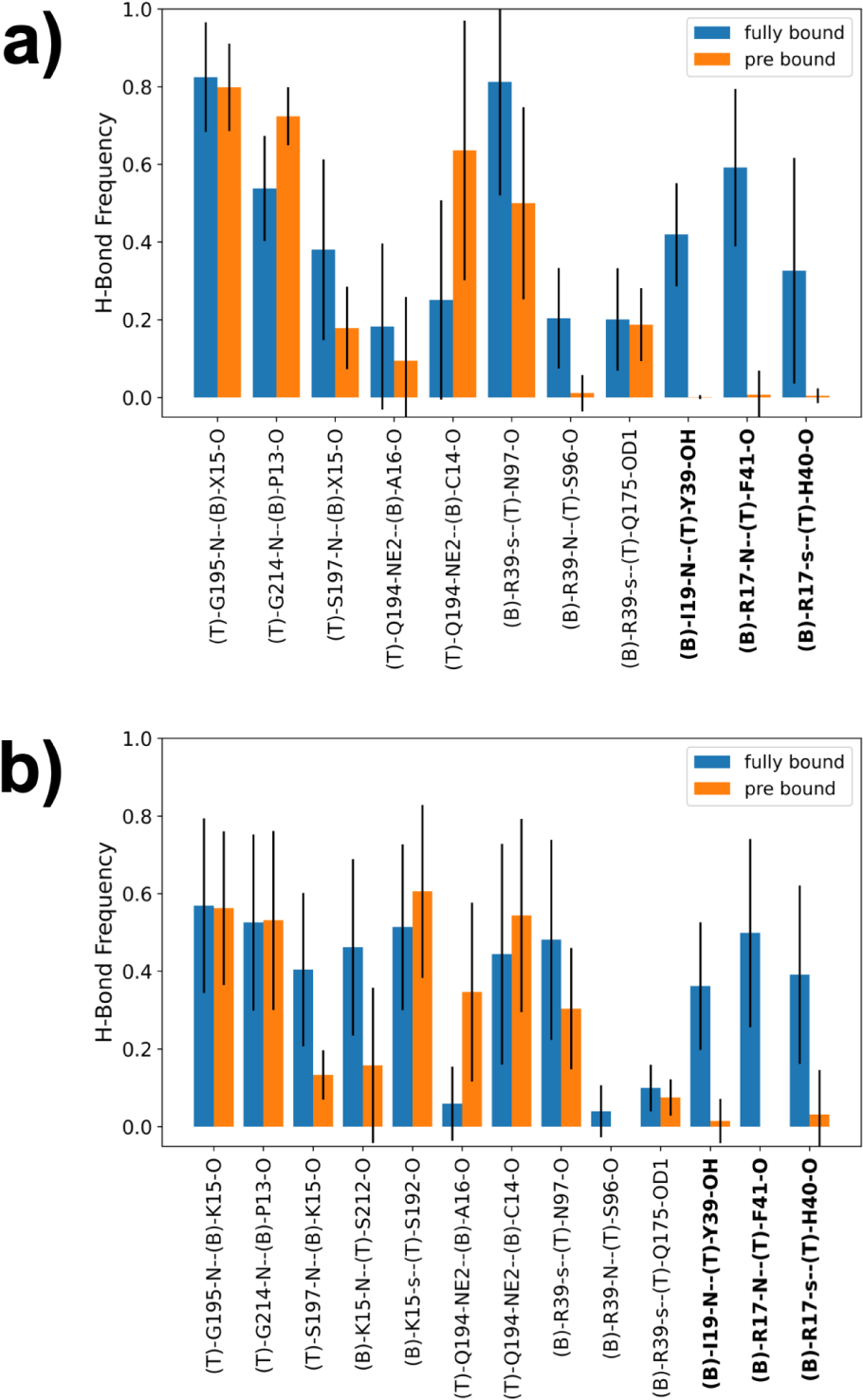
(a) Hydrogen bond frequencies of all combined unbiased simulations of the fully bound state and the pre-bound state. Residue name X = Abu, MfeGly, DfeGly or TfeGly, T = Trypsin, B = (Abu, MfeGly, DfeGly, TfeGly)-BPTI. O, N = heteroatoms in the backbone, OD1, OH, NE2 = heteroatoms in side chains. Hydrogen bonds are denoted as Donor-Acceptor. The side chain of arginine residues is denoted as “s”, which means a hydrogen bond with any of the donors in the guanidine moiety. (b) Hydrogen bond frequencies the unbiased simulations of the fully bound state and the pre-bound state with wildtype-BPTI. The labels follow the same scheme as above. The side chain of lysine is also denoted as “s”.

In the simulations of the fully bound state, we observe eleven hydrogen bonds with a relative population > 0.1. These are five hydrogen bonds, which are located around the S1 pocket, three hydrogen bonds of Arg39(compare fig.2b) and lastly the hydrogen bonds of Arg17 and Ile19 (compare fig.2c). We find the hydrogen bonds around the S1 pocket to also be present in the pre-bound state. The backbone-backbone interaction between Gly195 and the P1 residue has the same frequency in the pre-bound state as in the fully bound state, while the frequency of the neighboring interaction between the side chain of Gln194 and the backbone of Ala16 is lower in the pre-bound state, albeit with high statistical uncertainty. The frequency of two hydrogen bonds close to the S1 pocket, namely between the side chain of Gln194 and the backbone of Cys14 and the backbone-backbone interaction between Gly214 and Pro13 is higher in the pre-bound state, but again with high statistical uncertainty.

In the simulations of the complex with wildtype-BPTI, we find the same hydrogen bonds like for the Abu-BPTI variants (fig. 7b). Additionally, we observe a frequent hydrogen bond between the side chain of Lys15 and Ser192, which is a well known key interaction between trypsin and wildtype-BPTI at the bottom of the S1 pocket.^6^ As for the Abu-BPTI variants, the hydrogen bonds around the S1 pocket are in place in the fully bound state and in the pre-bound state. Interestingly, this also applies to the interaction between Lys15 and Ser192 at the bottom of the S1 pocket, meaning that in the pre-bound state, this key interaction of wildtype-BPTI is still in place.

Three hydrogen bonds are frequently populated in the fully bound state but are virtually non-existent in the pre-bound state, making these three broken hydrogen bonds a defining property of the pre-bound state. These are the same three hydrogen bonds that already showed a loss of population when transition from the fully bound state to the pre-bound state in the RAMD simulations (fig. 5.b). In the fully bound state, two of the hydrogen bonds are formed between between Arg17 in the BPTI variants and the backbone in trypsin, one between the side chain of Arg17 and the backbone of His40 and the other between the backbone of Arg17 and the backbone of Phe41. The third hydrogen bond is formed between the amide hydrogen of Ile19 in BPTI and the side chain of Tyr39 in trypsin. These hydrogen bonds are shown for the fully bound state in fig. 9.a. Fig. 9.b shows the same region in the pre-bound state. Side chains of Arg17 and Tyr39 have been reoriented and the three hydrogen bonds cannot be formed in the pre-bound state.

The hydrogen bonds of Arg39 in the BPTI variants with the backbone of trypsin are also more frequently populated in the fully bound state than in the pre-bound state (fig. 7). However, the drop in population is less pronounced than for the three hydrogen bonds discussed above. For wildtype-BPTI, the hydrogen bonds are less populated in the fully bound state and also in the pre-bound state, compared to the Abu-BPTI variants.

The analysis so far shows that the dissocation of the protein-protein complex between trypsin and (Abu, MfeGly, DfeGly, TfeGly)-BPTI proceeds via a pre-bound state which is stable at least on the timescale of 50 ns. The pre-bound state is characterized by a shift in the positional variables of BPTI and, to a lesser extent, by a shift in the orientational variables. To form the pre-bound state, three hydrogen bonds that are highly populated in the fully bound state, are broken.

### Stabilizing Interactions in the Pre-Bound State

The analysis so far does not show why the breaking of the three hydrogen bonds results in a stable state that does not immediately revert back to the fully bound state. Fig. 8.a suggests that one of the factors contributing to the stability of the pre-bound state could be a cation-pi interaction that is formed by the now free Arg17 side chain of BPTI with the aromatic system of Tyr151 of trypsin.

**Figure 8:**
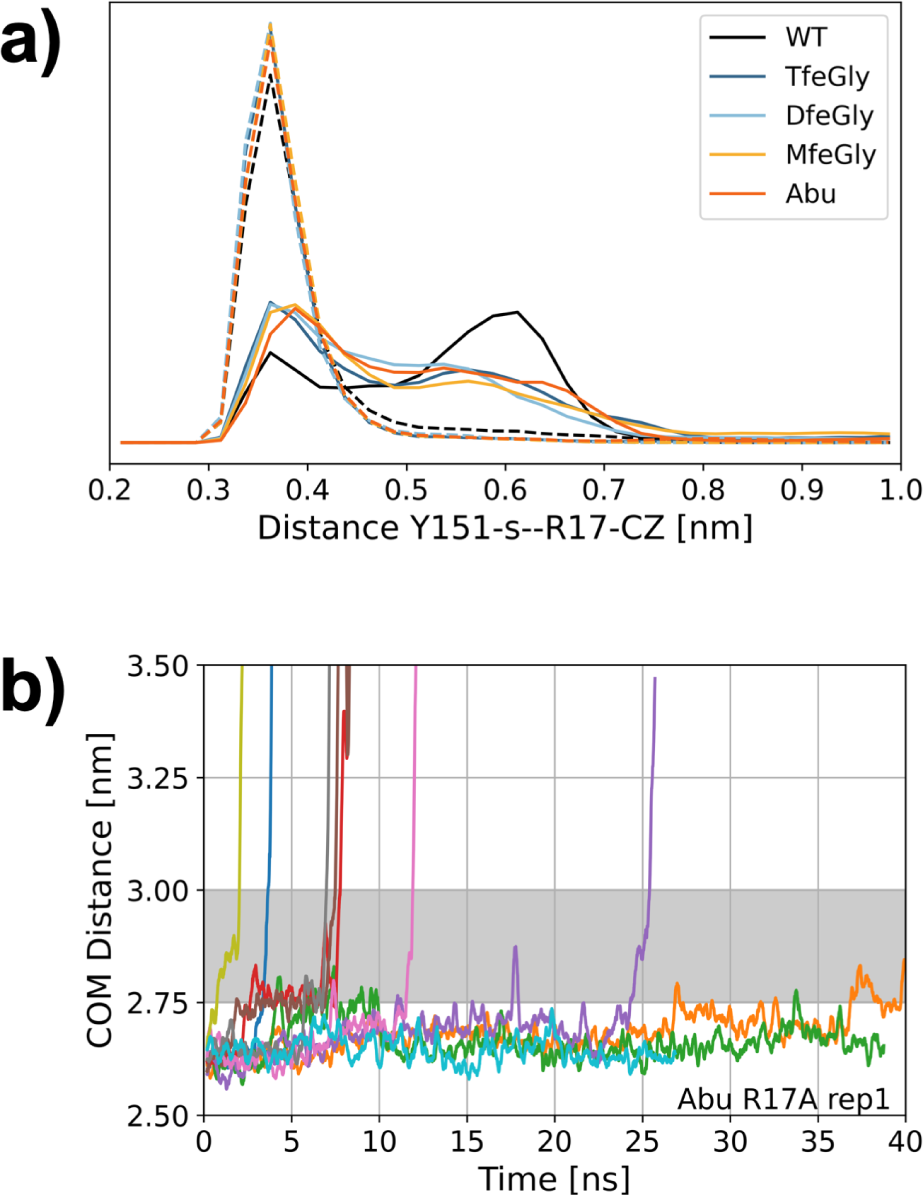
(a) Distance between the centroid of the Tyr151 aromatic system (Y151-s) and the carbon of the guanidine moiety of Arg17 (R17-CZ) in the unbiased simulations of the fully bound state (solid line) and the pre-bound state (dashed lines). (b) RAMD dissociation trajectories of one replica of the R17A mutant of Abu-BPTI.

We measured the distance distribution between the carbon atom of the guanidine moiety of Arg17 (Arg17-CZ) and the centroid of the aromatic ring of Tyr151 (fig.8a). While in the fully bound state, the distance can take a range of values between 0.3 nm and 0.8 nm, the distance in all simulation snapshots of the pre-bound state remains well below 0.45 nm. The broad distribution of the Tyr151-s-Arg17-CZ distance in the fully bound state shows that no specific bond is in place between the two residues. By contrast, the narrow distribution at low distances in the pre-bound state suggests the existence of a cation-pi interaction.

An interaction with the aromatic system of Tyr151 has so far not been described for the BPTI-trypsin complex. It is however present in X-ray crystallography structures of other trypsin inhibitors like Bdellastasin (pdb code: 1C9T), where a cationic lysine side chain at P2’ position forms a cation-pi interaction with Tyr151^46^ or Microviridin (pdb code: 4KTU) where a tyrosine at P2’ position forms a t-shaped pi-pi interaction with Tyr151.^47^

To verify whether an interaction of the Arg17 side chain is indeed essential for the stabilization of the pre-bound state, we repeated the RAMD simulations for one replica of the Abu-BPTI-trypsin complex, where we mutated Arg17 in Abu-BPTI to alanine (fig.8b). The dissociation happens roughly on the same timescale as for the non-mutated Abu-BPTI variants. However, the pre-bound state is traversed rapidly on all ten of the unbinding trajectories. This supports the hypothesis that Arg17 is indeed essential for the stabilization of the pre-bound state.

**Figure 9:**
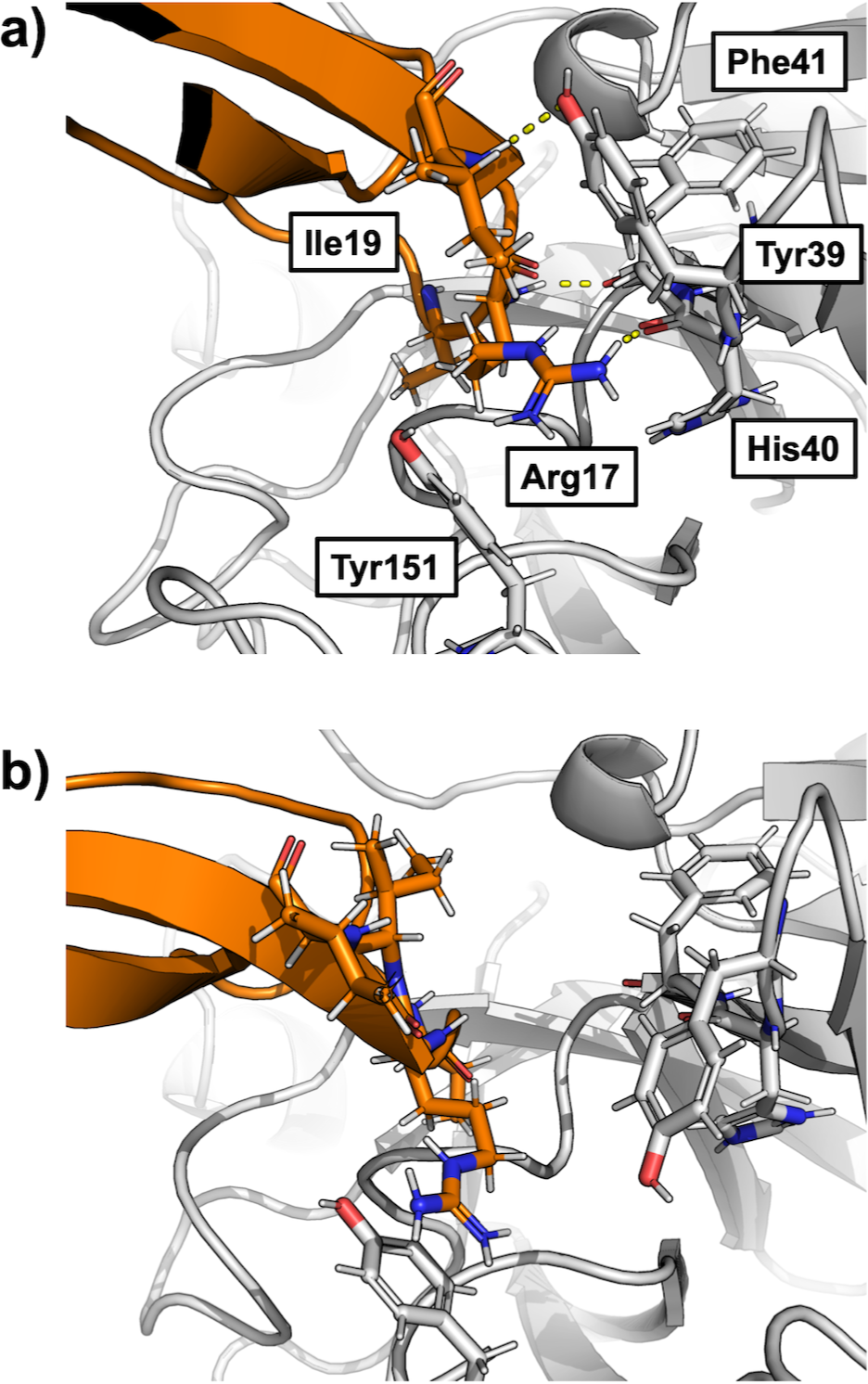
Example snapshots from the unbiased simulations of the (a) fully bound and (b) pre-bound state. The figure shows a similar region as fig. 2c.

Additionally, we analyzed the solvent accessible surface area (SASA) of the protein-protein interface amino acid residues for the fully bound state and pre-bound state (see fig. S14-S17). Most of the residues in the interface do not show significant differences in their SASA in the fully bound state and pre-bound state. A notable exception is that the SASA of the residues Arg17, Ile18 and Ile19 of the BPTI variants, as well as of Tyr39 and Phe41 of trypsin increases significantly in the pre-bound state. The SASA of Tyr151 decreases in the pre-bound state. These changes reflect the difference in binding between the fully bound state and the pre-bound state. This implies that the hydration shell of the fully bound state and the pre-bound state is similar, except for the region around Arg17. Thus, the pre-bound state is likely not only stabilized by the interaction of Arg17 and Tyr151, but other effects, such as hydration play a role as well.

### The Influence of the Fluorine Substituents

As a last step, we were interested in how the fluorine substituents in the BPTI variants influence the stability of the pre-bound state. To this end, we performed Umbrella Sampling between the fully bound state and the pre-bound state, where we used the newly identified interaction between the backbone amide of Arg17 in BPTI and the backbone oxygen of Phe41 in trypsin. We selected this reaction coordinate combined with a slow growth approach for the starting structures of the umbrella windows to ensure an accurate transition path between the fully bound state and the newly discovered pre-bound state. We find this approach to model the transition more accurately than picking starting structures from our RAMD simulations and using the center-of-mass distance as the reaction coordinate, as attempts to model the transition path using the string method with swarms-of-trajectories^48,49^ did not capture the transition state between the two states (see fig. S18).

Fig. 10a shows the resulting potential of mean force along this reaction coordinate derived from the newly identified interaction and the slow-growth approach. In all four systems, the potential of mean force exhibits two minima. The minimum around the a 0.3 nm corresponds to the fully bound state, whereas the minimum around 0.6 nm corresponds to the pre-bound state. In a previous study,^22^ we investigated the interactions in the fully bound state and found no significant differences between the four BPTI variants. For the pre-bound state, we find that the barrier height between the two states for the unfluorinated Abu and the mono-fluorinated MfeGly is about 15 kJ/mol, while for the higher fluorinated DfeGly and TfeGly it is only about 10 kJ/mol. The minimum of the pre-bound state for Abu lies well above the minimum for the fully bound state. By contrast, in the TfeGly-BPTI complex, the pre-bound state is stabilized relative to the bound state. The partially fluorinated complexes lie in between. Thus, there is a clear effect of the fluorination on the energetic landscape between the fully bound state and the pre-bound state.

**Figure 10:**
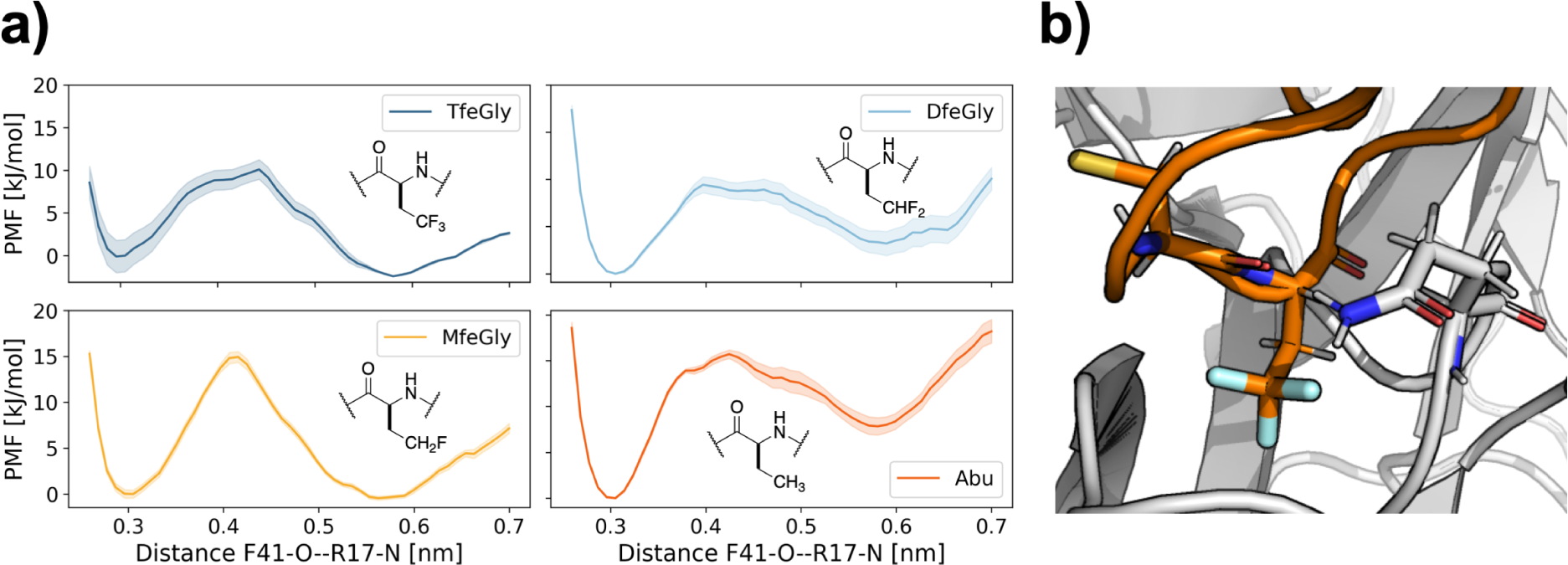
(a) Potential of mean force profile of fully bound state and pre-bound state from Umbrella Sampling over the distance between the carbonyl oxygen of Phe41 (F41-O) and the backbone nitrogen of Arg17 (R17-N). (b) Interaction between Gln194 and Cys14 in the direct proximity of the TfeGly side chain.

To find a possible mechanism for this stabilization, we revisited our hydrogen bond analysis for which we reported the aggregate statistics for all four Abu-BPTI variants in fig. 7a. We re-analyzed for each BPTI-variant and found that for most interactions, the hydrogen bond populations did not differ significantly across the BPTI variants. A notable exception is the hydrogen bond between the side chain of Gln194 in trypsin and the backbone oxygen of Cys14 in the BPTI variants. This interaction can be observed in the fully bound state and also in the pre-bound state, but it is more frequent in the pre-bound state of the fluorinated variants (MfeGly, DfeGly, TfeGly)-BPTI, while it is equally populated in the states of Abu-BPTI (see fig. S13). Gln194 and Cys14 are close to the side chain of the P1 residue (see fig.10b). When the hydrogen bond is formed, the side chain of Gln194 is in fact so close to the fluorine atoms that it appears plausible that the fluorine atoms with their negative partial charge help stabilize the NH2-end of the Gln194 side chain by providing an extra binding partner in addition to the backbone oxygen of Cys14.

## Conclusions

We applied several molecular simulation techniques to characterize the unbinding pathway of (Abu, MfeGly, DfeGly, TfeGly)-BPTI and wildtype-BPTI from trypsin. The BPTI variants likely dissociate via a curved pathway in a coordinate space that describes the relative position and orientation of the two proteins, as evidenced by restrained Metadynamics simulations in which the two protein do not rebind once they are dissociated. Using RAMD simulations^23–26^ to accommodate this curved unbinding pathway, we identified a new metastable state on the unbinding pathway.

This pre-bound state is present on the unbinding pathway in all four variants of the BPTI-trypsin complex and also in the wildtype-BPTI-trypsin complex. In unbiased simulations it is stable for at least 50 ns. Since in an aggregated simulation time of 1 *µ*s per BPTI variant, the pre-bound state only very rarely reverted to the fully bound state, we suspect that the average lifetime of the pre-bound state is in fact in the order of several 100 ns.

The pre-bound state is clearly distinct from the fully bound state in the positional coordinates from the fully bound state. The center-of-mass distance between the two proteins in the complexes of the Abu-BPTI variants is increased by about 0.2 nm (from 2.65 to 2.85 nm) and the BPTI variants rotate by about 10*^◦^* (0.2 rad) in Θ*_p_* and by 10*^◦^* (0.2 rad) around the dihedral angle Φ*_p_*. There is little overlap between the distributions of the pre-bound state and fully bound state in these coordinates. We also observe a systematic shift in the orientational coordinates, but less pronounced. The distribution of fully bound state and pre-bound state for wildtype-BPTI is very similar to those of the Abu-BPTI variants, with the exception of Θ*_p_*, which is slighly shifted towards higher values.

The interaction pattern between the two proteins changes when transitioning from the fully bound state to the pre-bound state. These changes particularly involve Arg17 (P2’ residue) and Arg39 in the BPTI variants. In the pre-bound state, the hydrogen bond of the Arg17 side chain to the backbone of trypsin is broken, but it is replaced by a cation-pi interaction between the guanidine moiety and a nearby trypsin tyrosine residue. Two further hydrogen bonds in the vicinity are also broken in this process, and the hydrogen bond between the side chain of Arg39 and the trypsin backbone becomes less populated. When we replaced Arg17 by an alanine residue in RAMD simulations, the protein-protein complex dissociated without spending time in the pre-bound state, which demonstrates that Arg17 is essential for the stabilization of this state.

The pre-bound state is likely not only stabilized by the interaction of Arg17 and Tyr151, but also due to other effects, such as hydration. The SASA is increased for the residues close to Arg17 in the pre-bound state, which might imply a change in hydration. This aspect should be addressed in future research, e.g. by an analysis of the water molecules in the vicinity of Arg17 similar to our analysis of the water molecules in the S1 binding pocket. ^22^

The structural rearrangements that stabilize the pre-bound state do not involve the P1 residue in BPTI or the negatively charged residues at the bottom of the S1 pocket of trypsin. The same structural rearrangements can also be found for wildtype-BPTI, which means the unbinding of the Abu-BPTI variants proceed via the same pre-bound state.

In potentials of mean force (PMF), we find that fluorination of Abu lowers the free energy barrier between the fully bound and the pre-bound state and also lowers the free energy minimum of the pre-bound state. However, a quantitative interpretation of these one-dimensional PMFs is difficult. In particular, we suspect that the PMF might overstabilize the pre-bound state, as in some of the potentials the pre-bound state minimum is as low as the fully bound minimum. Nonetheless, the fluorine substituents on the P1 residue clearly have an influence on the stability of the pre-bound state. A possible, yet speculative, explanation is that the hydrogen bond between the side chain of Gln194 and Cys14 is stabilized by fluorine substituents in direct proximity of the side chain NH2 group of Gln194. Fluorine is known to have a wide range of possible effects on protein-inhibitor interactions, e.g. through hydrogen bonds,^50^ desolvation^33,34^ or entropy,^51^ whose elucidation often requires in-depth computational studies. The differences in barrier height and stability of the pre-bound state in the fluorinated variants of BPTI are likely not only due to a single stabilizing interaction, like the Gln194-Cys14 hydrogen bond, but instead due to a combination of enthalpic and entropic effects.

Because of the large magnitude of the biasing force in the RAMD simulations, which is necessary to dissociate the protein-protein complexes, we did not observe encounter complexes in our simulations. We expect that encounter states do play a role in the binding and unbinding process of the (Abu, MfeGly, DfeGly, TfeGly)-BPTI-trypsin complexes.^20^ However, the transition between the pre-bound state and these weakly bound encounter states should be characterized with other methods like Weighted Ensemble MD^52^ and Molecular Rotational Grids.^53^

While this manuscript was in review, D’Arrigo et al. ^54^ published a preprint, in which they dissociate a series of protein-protein systems, including wildtype-BPTI and some of its mutants from trypsin, using RAMD with a smaller force. In the dissociation trajectories, they find that the contacts of Arg17 are cleaved first, which aligns well with our results. The authors find additional states along the dissociation trajectory, which may correspond to the encounter states mentioned above. These additional states, together with works of Kahler et al.^20^ are excellent starting points for the characterization of encounter states that we suggest above.

The existence and structure of the pre-bound state invites speculation on the inhibitory mechanism of BPTI and its variants. After forming the initial Michaelis complex of a substrate with trypsin, the hydrolysis of the peptide bond proceeds via two steps. First the peptide bond is broken and the N-terminal part of the substrate (i.e., all residues from the N-terminus up to and including P1) forms a covalently bound acyl-enzyme intermediate. The C-terminal part of the substrate (i.e., all residues from P1’ to the C-terminus) remains non-covalently bound and needs to dissociate before, in a second step, the acyl-enzyme intermediate can be hydrolyzed. Radisky and Koshland showed that, for a closely related serine protease complex, the initial formation of the acyl-enzyme intermediate is fast, but the release of the C-terminal part of the substrate is slow, ^9^ such that the reaction reverts back to the intact peptide bond. This “clogged gutter” mechanism is further supported by a high-resolution structure of a cleaved BPTI variant with trypsin.^13^ Our analysis showed that the interface between trypsin and the BPTI variants is stabilized by hydrogen bonds primarily from the C-terminal part of the BPTI variants (fig.7). Specifically, Arg17 which stabilizes the pre-bound state via a cation-pi interaction belongs to the C-terminal part. Thus, assuming that the clogged gutter mechanism applies to the BPTI-trypsin complex, these interactions likely contribute to stabilizing the C-terminal part within the protein complex.

Finally, our study shows that, to understand the stability of the wildtype-BPTI-trypsin complex or the (Abu, MfeGly, DfeGly, TfeGly)-BPTI-trypsin complex, one needs to consider two states, the fully bound state and the pre-bound state, which likely are in dynamic equilibrium. By mimicking the interactions in the pre-bound state one may open up additional ways to design serine-protease inhibitors.

## Supporting information

Example Snapshots

## Data And Software Availability

RAMD and unbiased MD simulations were performed with openly available Gromacs (https://www.gromac Gromacs-RAMD (https://github.com/HITS-MCM/gromacs-ramd/tree/release-2022) and Plumed (https://www.plumed.org) software. Starting structure preparation was done with the openly available pdbfixer (https://github.com/openmm/pdbfixer). Interaction distances and hydrogen bonds were analyzed using mdtraj (https://www.mdtraj.org), which is openly available. Force field parameters for the fluorinated amino acids and Abu have been released in a recent publication,^22^ all other force field parameters were taken from the openly available Amber14SB force field (https://ambermd.org). Protein structure starting files, MD parameter files and analysis scripts for all simulations are published on Github (https://github.com/leonwehrhan/ Trypsin BPTI Prebound RAMD 2024).

## Acknowledgement

We thank Ariane Nunes-Alves for fruitful discussions on RAMD simulations. Funded by the Deutsche Forschungsgemeinschaft (DFG, German Research Foundation): Project-ID 387284271-SFB 1349. The authors acknowledge access to high-performance computers via the Zentraleinrichtung FUB-IT of Freie Universität Berlin.

## Supporting Information Available

Center-of-mass distance timeseries for restrained Metadynamics simulations; Center-of-mass distance timeseries for RAMD simulations of TfeGly-, DfeGly-, Mfegly-, Abu-BPTI and wildtype BPTI; Positional and orientational collective variable histograms for the RAMD simulations; Center-of-mass distance timeseries for unbiased MD simulations; Orientational collective variable histograms for the unbiased MD simulations; Hydrogen bond frequencies for TfeGly-, DfeGly-, Mfegly-, Abu-BPTI and wildtype BPTI simulations. SASA for protein-protein interface residues of TfeGly-, DfeGly-, Mfegly-, Abu-BPTI. RAMD dissociation trajectories with higher forces. Example snapshots of the fully bound state and the pre-bound state of the four variants of the (TfeGly, DFeGly, MfeGly, Abu)-BPTI-trypsin complex.

## TOC Graphic

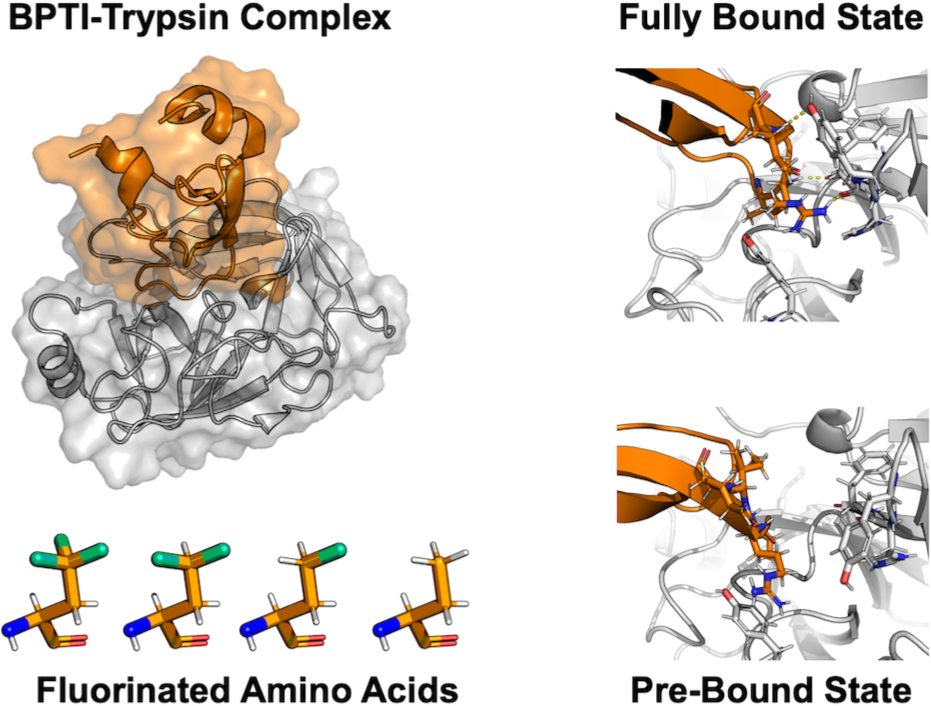

