## Supporting Information for "Pre-bound State Discovered in the Unbinding Pathway of Fluorinated Variants of the Trypsin-BPTI Complex Using Random Acceleration Molecular Dynamics Simulations"

### I. RESTRAINED METADYNAMICS

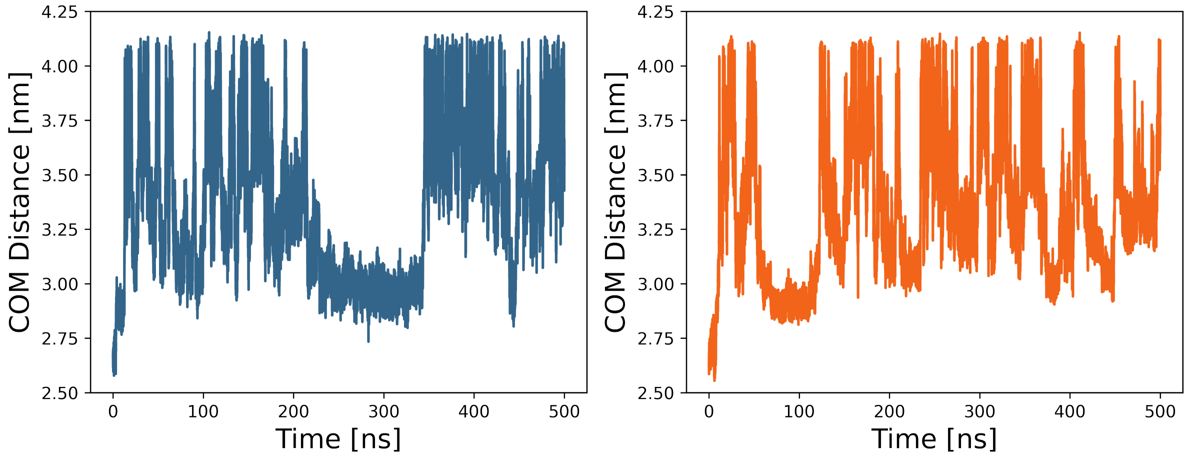


Figure S1. Center-of-mass (COM) distance between trypsin and BPTI as P1 TfeGly variant (left) and P1 Abu variant (right) throughout Metadynamics simulations with positional restraints.

#### A. Computational Methods

*a. Restrained Metadynamics.* Metadynamics simulations were run with Gromacs 2021.5 and Plumed 2.8. Well tempered Metadynamics simulations were employed for studying the undinding process of BPTI from Trypsin. The collective variable was defined to be the backbone center of mass distance between the receptor trypsin and the ligand BPTI. The repulsive gaussians were deposited at a rate of 1 ps at an initial height of 1.2 kJ/mol and $\sigma$ of 0.1 nm. The biasfactor was set to be 5. An upper wall at a center of mass distance of 4.0 nm was installed with a force constant of 2500 kJ/(mol$\cdot$nm$^2$). The collective variables $\Theta_p$, $\Phi_p$, $\theta_1$, $\phi_1$ and $\psi_1$ were constrained with harmonic potentials of 400 kJ/(mol $\cdot$ rad$^2$) to remain at their respective value in the crystal structure of $\beta$-Trypsin-TfeGly-BPTI (pdb code: 4Y11). The orientational angle $\theta_1$ and the orientational dihedrals $\phi_1$ and $\psi_1$ were restrained at values of 2.068 rad, -0.956 rad and 0.592 rad, respectively. The positional dihedrals $\Theta_p$ and $\Phi_p$ were restrained at 1.991 rad and 1.437 rad, respectively.

### II. RAMD TRAJECTORIES

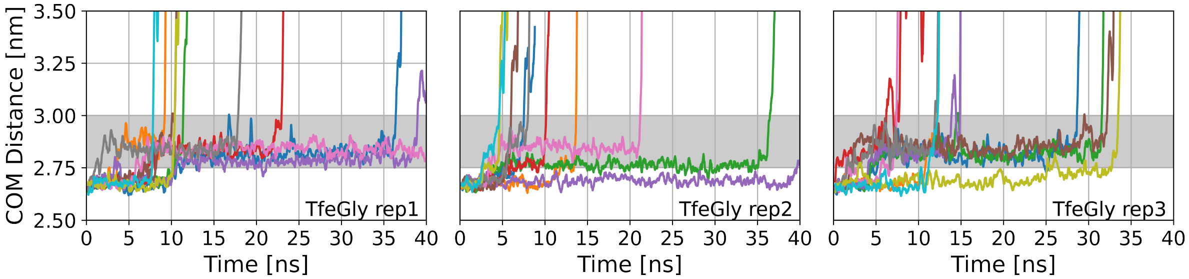


Figure S2. RAMD dissociation timeseries of TfeGly-BPTI. COM = center-of-mass. Grey area shows the center-of-mass distance of the pre-bound state. Left panel: replica 1, middle panel: replica 2, right panel:replica 3. 10 RAMD runs per replica.

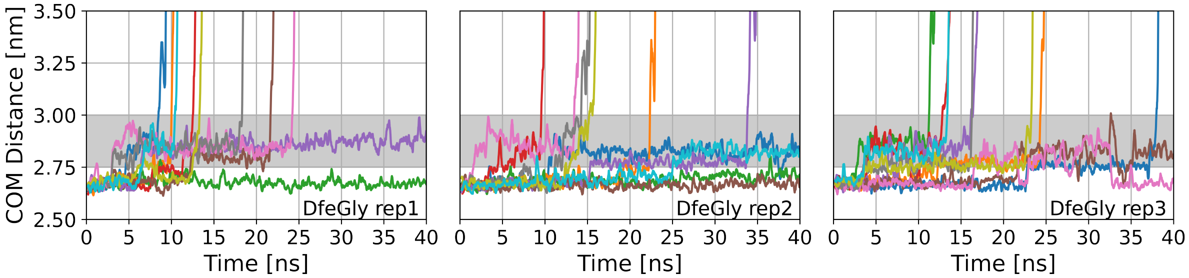


Figure S3. RAMD dissociation timeseries of DfeGly-BPTI. COM = center-of-mass. Grey area shows the center-of-mass distance of the pre-bound state. Left panel: replica 1, middle panel: replica 2, right panel:replica 3. 10 RAMD runs per replica.

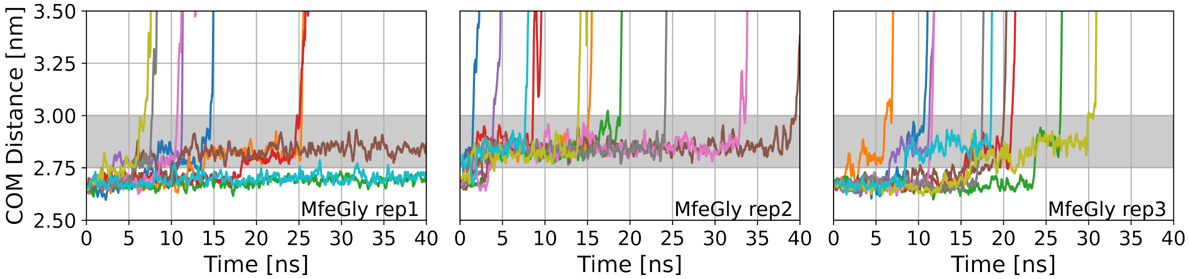


Figure S4. RAMD dissociation timeseries of MfeGly-BPTI. COM = center-of-mass. Grey area shows the center-of-mass distance of the pre-bound state. Left panel: replica 1, middle panel: replica 2, right panel:replica 3. 10 RAMD runs per replica.

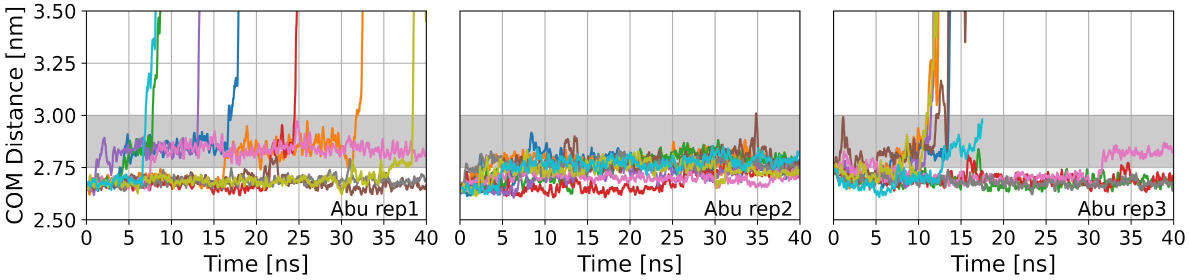


Figure S5. RAMD dissociation timeseries of Abu-BPTI. COM = center-of-mass. Grey area shows the center-of-mass distance of the pre-bound state. Left panel: replica 1, middle panel: replica 2, right panel:replica 3. 10 RAMD runs per replica.

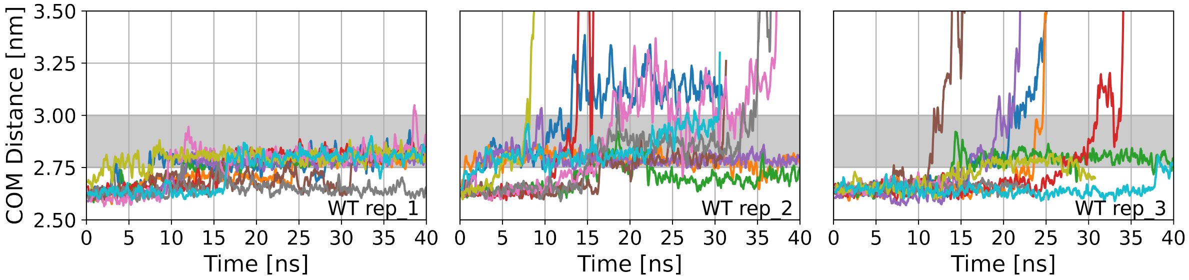


Figure S6. RAMD dissociation timeseries of wildtype-BPTI. COM = center-of-mass. Grey area shows the center-of-mass distance of the pre-bound state. Left panel: replica 1, middle panel: replica 2, right panel:replica 3. 10 RAMD runs per replica.

TABLE S1. RAMD simulations of complexes of trypsin with (fluorinated) Abu-BPTI complexes. For every replica there are 10 simulations of max. 40 ns length, which only differ by the random seed for the RAMD force. The criterion for the pre-bound state to be observed is that, throughout the whole simulation, the 200 ps moving average of the center-of-mass distance has to be at least 1 ns without interruption between 2.75 nm and 3.00 nm.

| Complex | Replica | Dissociated | Undissociated | Pre-bound observed |
|---|---|---|---|---|
| TfeGly | 1 | 8 | 2 | 8 |
| TfeGly | 2 | 9 | 1 | 6 |
| TfeGly | 3 | 10 | 0 | 8 |
| DfeGly | 1 | 8 | 2 | 7 |
| DfeGly | 2 | 6 | 4 | 7 |
| DfeGly | 3 | 8 | 2 | 10 |
| MfeGly | 1 | 7 | 2 | 7 |
| MfeGly | 2 | 9 | 1 | 9 |
| MfeGly | 3 | 10 | 0 | 7 |
| Abu | 1 | 7 | 3 | 7 |
| Abu | 2 | 0 | 10 | 7 |
| Abu | 3 | 6 | 4 | 4 |

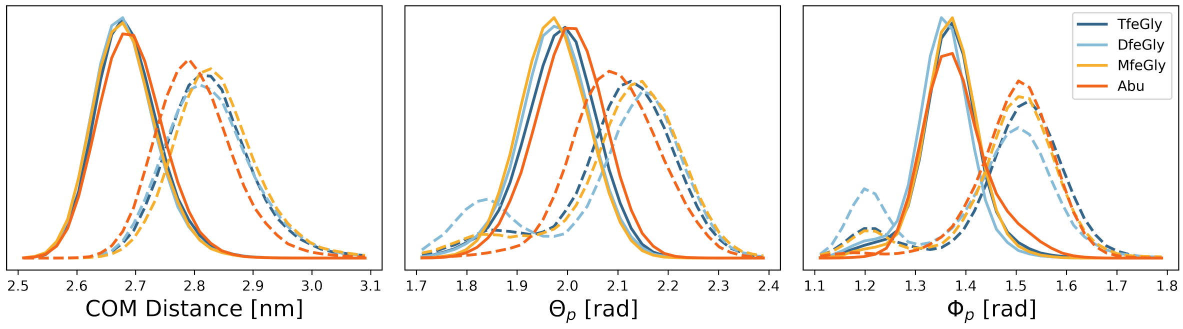


Figure S7. Center-of-mass distance histogram (left) histograms of $\Theta_p$ (middle) and $\Phi_p$ (right) for the fully bound state (solid line) and pre-bound state (dashed line) in the RAMD simulations of the Abu-BPTI variants.

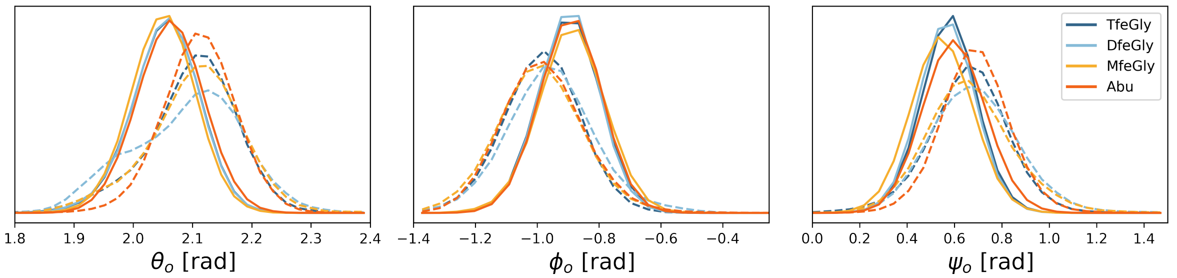


Figure S8. Histograms of $\theta_o$ (left), $\phi_o$ (middle) and $\psi_o$ (right) for the fully bound state (solid line) and pre-bound state (dashed line) in the RAMD simulations of the Abu-BPTI variants.

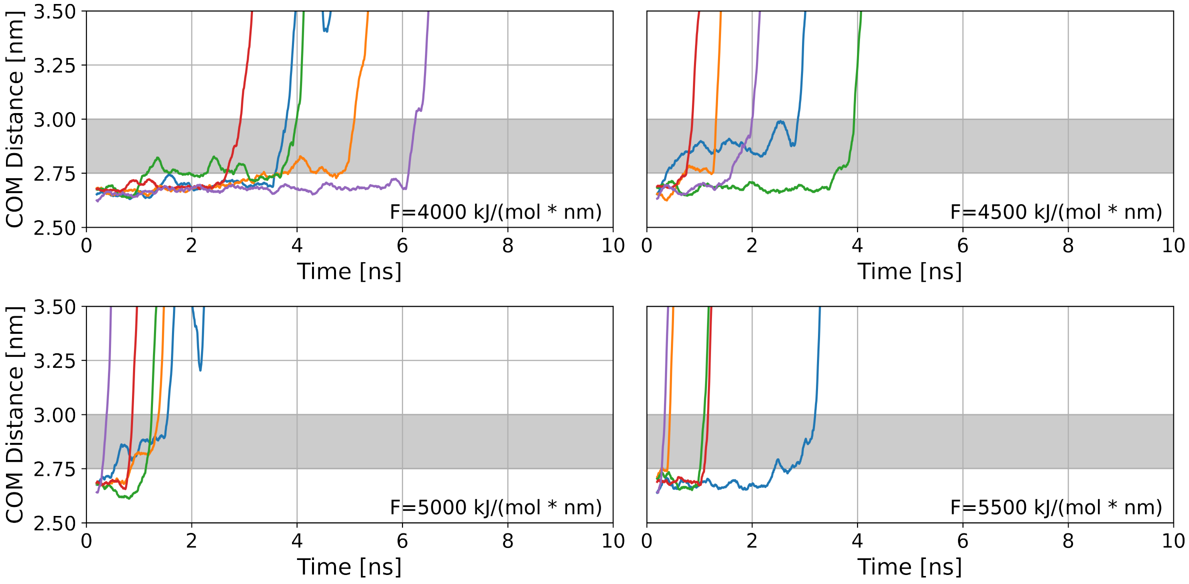


Figure S9. RAMD dissociation trajectories of TfeGly-BPTI with high random forces.

### III. UNBIASED SIMULATIONS

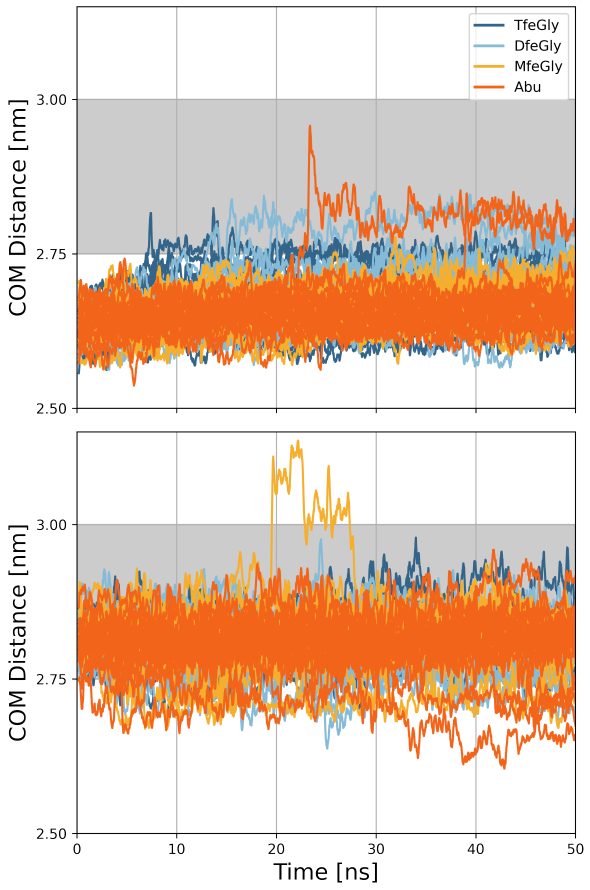


Figure S10. COM distance trajectory of all RAMD simulations of the Abu-BPTI variants overlayed. The grey area indicates the pre-bound state. The fluctuations throughout the trajectory are smoothed by calculating a moving average with a moving window of 200 ps.

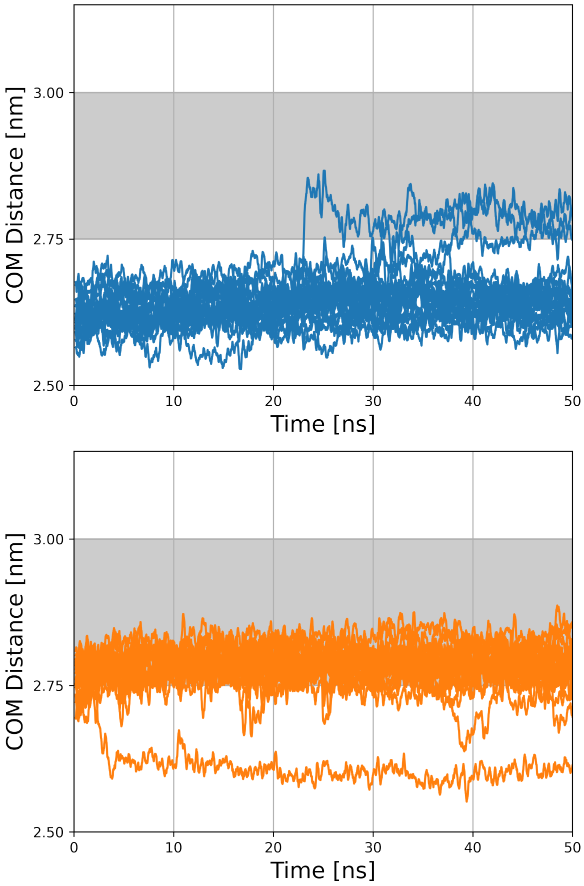


Figure S11. COM distance trajectory of all RAMD simulations of wildtype-BPTI overlayed. The grey area indicates the pre-bound state. The fluctuations throughout the trajectory are smoothed by calculating a moving average with a moving window of 200 ps. The simulations in the top panel were started in the fully bound state, the simulations in the bottom panel were started in the pre-bound state.

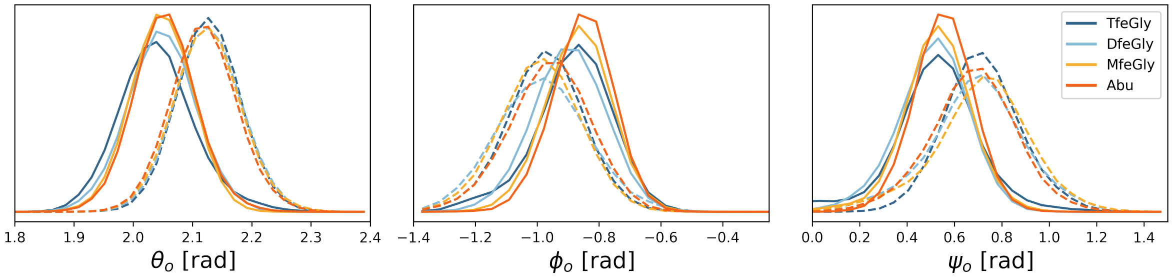


Figure S12. Histograms of $\theta_o$ (left), $\phi_o$ (middle) and $\psi_o$ (right) for the fully bound state (solid line) and pre-bound state (dashed line) in the unbiased simulations of the Abu-BPTI variants.

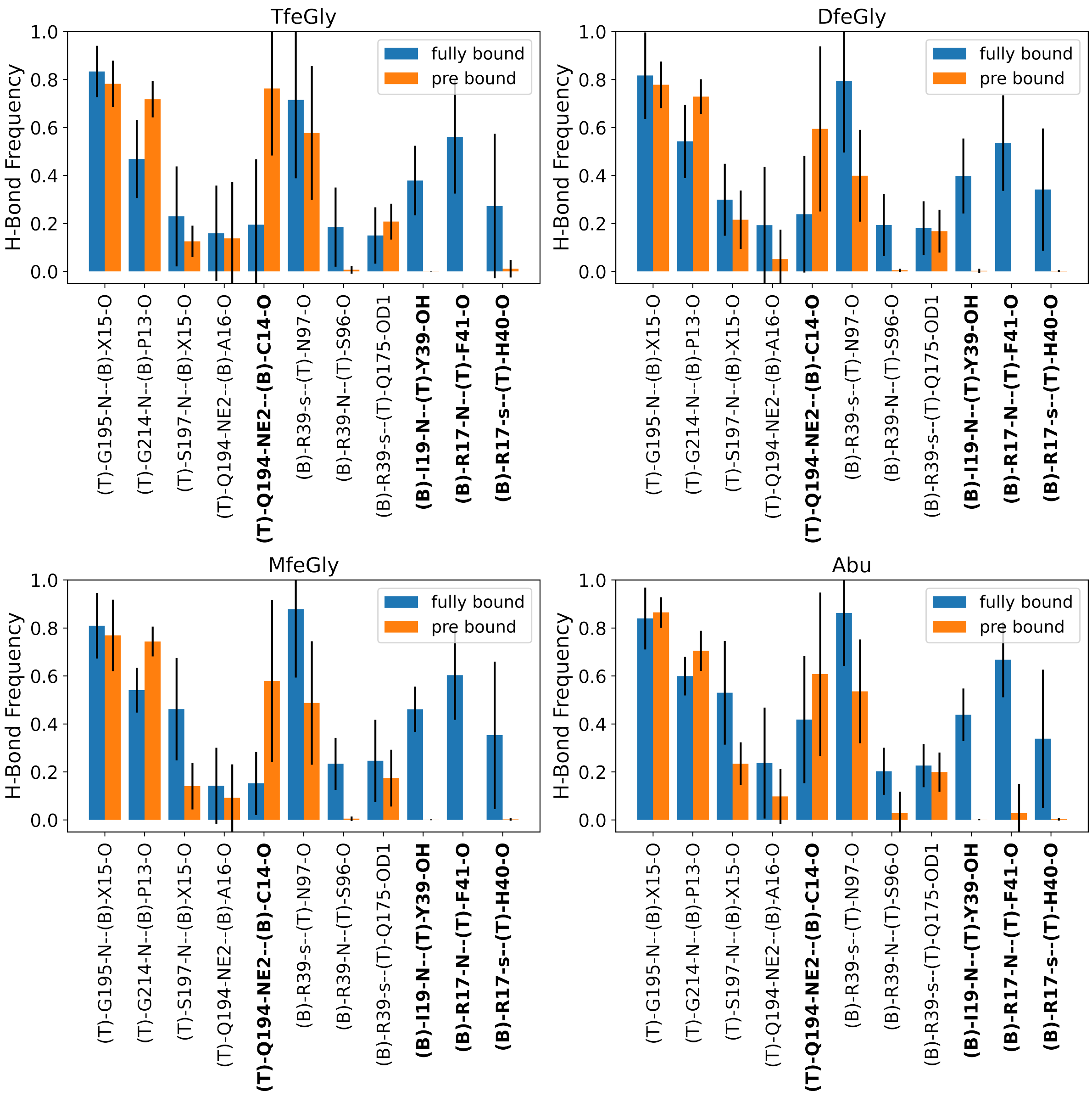


Figure S13. Hydrogen bond frequencies of all the unbiased simulations of the fully bound state and the pre-bound state, separate for the four Abu-BPTI variants. Residue name X = Abu, MfeGly, DfeGly or TfeGly, T = Trypsin, B = (Abu, MfeGly, DfeGly, TfeGly)-BPTI. O, N = heteroatoms in the backbone, OD1, OH, NE2 = heteroatoms in side chains. Hydrogen bonds are denoted as Donor- Acceptor. The side chain of arginine residues is denoted as ”s”, which means a hydrogen bond with any of the donors in the guanidine moiety.

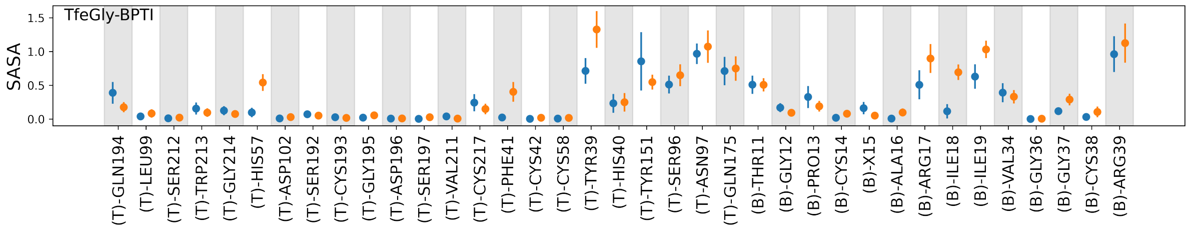


Figure S14. Solvent Accessible Surface Area (SASA) of all residues of the protein-Protein interface of trypsin and TfeGly-BPTI. (T) = trypsin, (B) = BPTI, X = TfeGly.

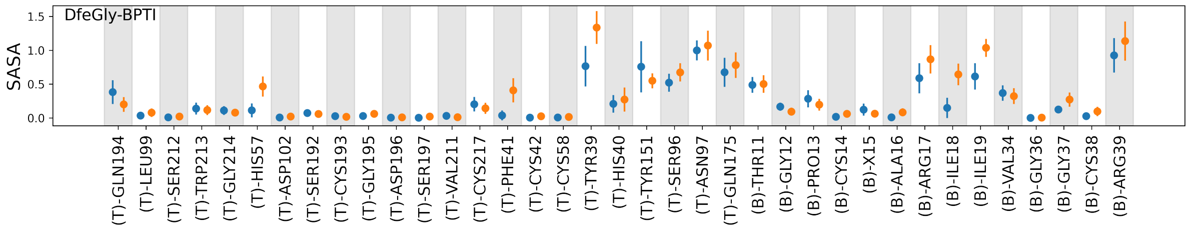


Figure S15. Solvent Accessible Surface Area (SASA) of all residues of the protein-Protein interface of trypsin and DfeGly-BPTI. (T) = trypsin, (B) = BPTI, X = DfeGly

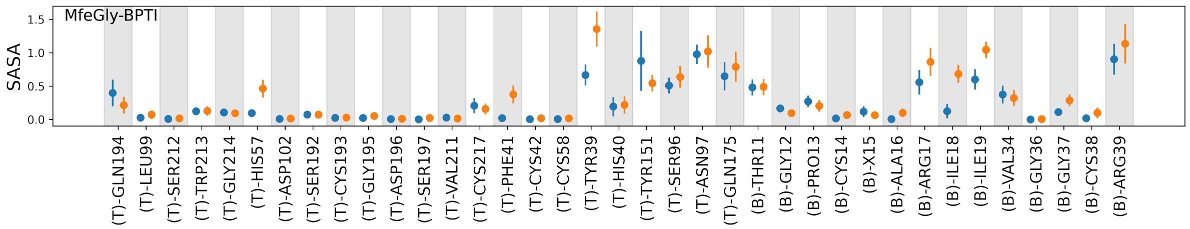


Figure S16. Solvent Accessible Surface Area (SASA) of all residues of the protein-Protein interface of trypsin and MfeGly-BPTI. (T) = trypsin, (B) = BPTI, X = MfeGly

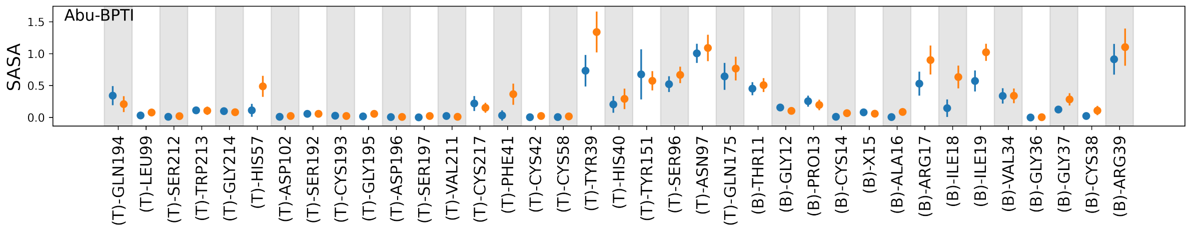


Figure S17. Solvent Accessible Surface Area (SASA) of all residues of the protein-Protein interface of trypsin and Abu-BPTI. (T) = trypsin, (B) = BPTI, X = Abu

### IV. SHORT EQUILIBRATION OF RAMD SNAPSHOTS FOR SWARMS-OF-TRAJECTORIES STRING METHOD

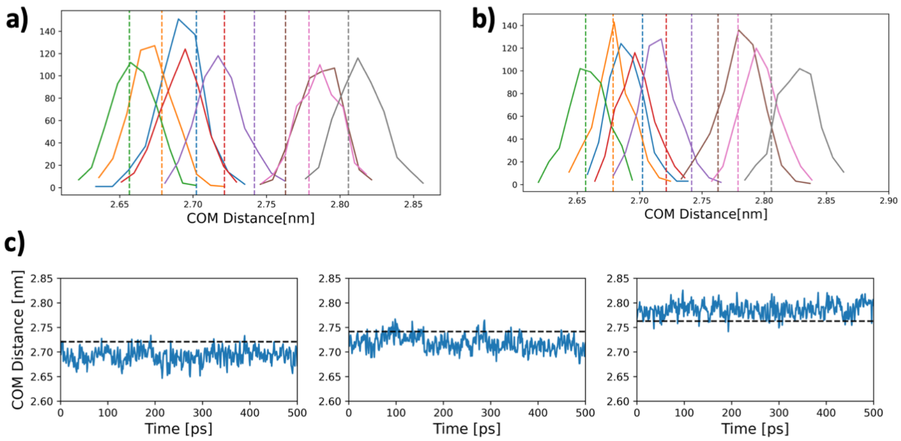


Figure S18. Short equilibration trajectories in our attempts for the swarms-of-trajectories string method. (a) Histograms of the distribution of the equilibration trajectories along the center-of-mass distance. The simulations are restrained on $r$, $\Theta_p$ and $\Phi_p$. $k = 6500kJ/mol/nm^2$ for $r$ and $k = 400kJ/mol/\text{rad}^2$ for $\Theta_p$ and $\Phi_p$. The equilibrium value of the restraint = initial value along $r$ is indicated as dashed line. (b) Histograms of the distribution of the equilibration trajectories along the center-of-mass distance. The simulations are restrained on the distance between Phe41-O and Arg17-N. $k = 6500kJ/mol/nm^2$ (c) Trajectory timeseries of the simulations seen in (a). The equilibrium value of the restraint = initial value is indicated as dashed line.
